# Protein embeddings improve phage-host interaction prediction

**DOI:** 10.1101/2023.02.26.530154

**Authors:** Mark Edward M. Gonzales, Jennifer C. Ureta, Anish M.S. Shrestha

## Abstract

With the growing interest in using phages to combat antimicrobial resistance, computational methods for predicting phage-host interactions have been explored to help shortlist candidate phages. Most existing models consider entire proteomes and rely on manual feature engineering, which poses difficulty in selecting the most informative sequence properties to serve as input to the model. In this paper, we framed phage-host interaction prediction as a multiclass classification problem, which takes as input the embeddings of a phage’s receptor-binding proteins, which are known to be the key machinery for host recognition, and predicts the host genus. We explored different protein language models to automatically encode these protein sequences into dense embeddings without the need for additional alignment or structural information. We show that the use of embeddings of receptor-binding proteins presents improvements over handcrafted genomic and protein sequence features. The highest performance was obtained using the transformer-based protein language model ProtT5, resulting in a 3% to 4% increase of weighted F1 scores across different prediction confidence threshold,compared to using selected handcrafted sequence features.

**Author summary:** Antimicrobial resistance is among the major global health issues at present. As alternatives to the usual antibiotics, drug formulations based on phages (bacteria-infecting viruses) have received increased interest, as phages are known to attack only a narrow range of bacterial hosts and antagonize the target pathogen with minimal side effects. The screening of candidate phages has recently been facilitated through the use of machine learning models for inferring phage-host pairs. The performance of these models relies heavily on the transformation of raw biological sequences into a collection of numerical features. However, since a wide array of potentially informative features can be extracted from sequences, selecting the most relevant ones is challenging. Our approach eliminates the need for this manual feature engineering by employing protein language models to automatically generate numerical representations for specific subsets of tail proteins known as receptor-binding proteins. These proteins are responsible for a phage’s initial contact with the host bacterium and are thus regarded as important determinants of host specificity. Our results show that this approach presents improvements over using handcrafted genomic and protein sequence features in predicting phage-host interaction.

## Introduction

One of the most pressing threats to global health is antimicrobial resistance (AMR), a phenomenon wherein microorganisms evolve to withstand exposure to bacteriostatic and bactericidal drugs. In 2019, 4.95 million AMR-related and 1.27 million AMR-attributable deaths were estimated [1]. In developing countries, this problem is compounded by the unregulated dispensation of antibiotics as a form of self-medication even for mild conditions [2, 3] and their routine use in the agricultural sector for disease prophylaxis [4] and livestock growth promotion [5].

A solution that is actively being explored to combat this problem is phage therapy, which capitalizes on the specificity of bacteriophages (hereinafter referred to as *phages*) to a narrow range of hosts. Phages have been shown to antagonize the target bacteria with minimal side effects and without triggering a dysbiosis of the beneficial microbiota [6]. However, the foremost challenge to formulating phage cocktails for treating bacterial infections is identifying putative phages that attack the offending pathogens. Aside from being time- and cost-intensive, *in vitro* experiments require the cultivation of microbes under strict laboratory conditions, posing a bottleneck to the rapid selection of candidate phages.

With the advent of high-throughput sequencing technologies and the resulting increase in omic data, *in silico* approaches have been employed to help shortlist candidate phages. These can be broadly categorized into alignment-based methods [7, 8], which rely on sequence similarity to infer phage-host pairs, and alignment-free methods [9–11], which exploit features related to sequence composition, such as oligonucleotide frequency and codon usage bias. These reflect shared genomic properties that arise from the close coexistence and coevolution of phages and their hosts [12, 13].

Machine learning algorithms for phage-host interaction prediction have also been actively explored. Feature sets extracted from protein sequences include molecular weight [14], aromaticity [15], amino acid composition [14], protein-protein and domain-domain interaction [14, 16], and protein secondary structure [15]. Meanwhile, those obtained from genomic sequences include *k*-mer frequency [14, 15, 17], guanine-cytosine content [15], codon usage bias [15, 16], oligonucleotide frequency [16, 18], and shared transfer ribonucleic acids [19]. Recent studies have also investigated the application of deep learning architectures, primarily convolutional neural networks, that take these handcrafted properties as input [20–23].

While these existing models have been successful in integrating various features to improve their performance, most consider the entire proteome of both the phages and their hosts [14, 16, 20, 21], when only specific proteins are actually involved in phage-host interaction [24]. To initiate infection, a tailed phage typically adsorbs to the host bacterium’s surface through receptor-binding proteins (RBPs) located at its tail’s distal end [25]. In this regard, RBPs (e.g., tail fibers and spikes) serve as key machinery for host recognition and specificity [26–29].

Moreover, most tools for phage-host interaction prediction rely on manual feature engineering to transform raw sequences into numerical vectors, often requiring additional alignment or structural information [30–32]. The multitude of potentially informative signals that can be derived from these sequences also poses difficulty in selecting which features should be fed to the model [30].

Representation learning, in which raw biological sequences are converted into dense vectors in high-dimensional space, has recently been applied to some prototypical bioinformatics tasks, such as predicting protein function [33], succinylation sites [34], and sequence conservation [35]. The embeddings produced by protein language models, such as SeqVec [36] (which adopts the architecture of the natural language model ELMo [37]) and the transformer-based Evolutionary Scale Modeling (ESM) [38] and ProtTrans [31], have been demonstrated to capture protein secondary structure and physicochemical characteristics, features that are relevant to phage-host specificity [31, 38]. Representation learning also serves as a promising approach to address some of the limitations posed by the problem of data scarcity in phage-host datasets [13], as knowledge is distilled from large-scale protein databases [39–42]. However, application of protein embeddings to the problem of phage-host interaction prediction remains unexplored.

In an attempt to address these gaps, our study seeks to contribute the following:

- We framed phage-host interaction prediction as a multiclass classification problem, with the embedding of a receptor-binding protein (RBP) as the input and the host genus as the output.
- We extensively tested different protein language models to automatically generate dense embeddings of RBP sequences.
- We constructed a random forest model for predicting phage-host interaction and showed that embeddings of RBPs outperform handcrafted genomic and protein sequence features, with the use of the protein language model ProtT5 resulting in the highest performance.

## Materials and methods

### Data collection and preprocessing

We collected genome sequences of 18,389 phages, along with their proteome sequences (whenever available), via INPHARED [43], a pipeline for retrieving phage sequences from GenBank [44]; the sequences were downloaded in September 2022, and proteomes were retrieved for 16,836 phages. Limiting the host information to the genus level, we subjected the entries to preprocessing of host information and selection of annotated receptor-binding proteins, which we describe in detail in the following subsections and summarize alongside our methodology in Fig 1.

**Fig 1.**
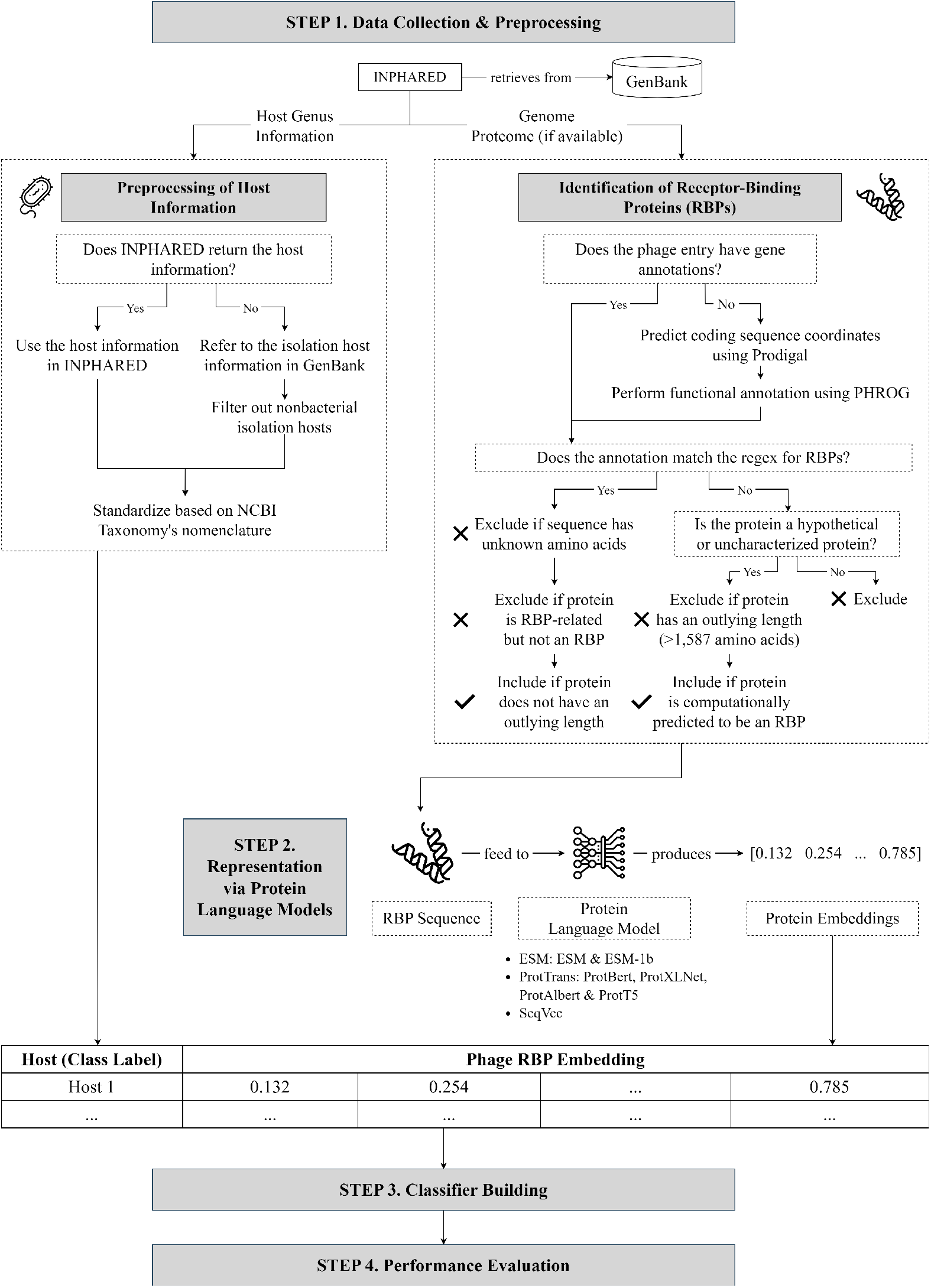
Methodology of our study. First, we collected phage genomes, along with their proteomes (whenever available), via INPHARED [43] and performed preprocessing to obtain the host information and select annotated receptor-binding proteins (RBPs). Second, we fed the RBP sequences to pretrained protein language models to generate meaningful dense embeddings. Third, we built a random forest model with the RBP embeddings as the input and the host genus as the predicted output. Finally, we evaluated our model’s performance. Flat icons used in this figure are taken from [45–47].

#### Preprocessing of host information

INPHARED [43] returns host data for 15,739 phages across 278 different host genera. For entries where the host name is unspecified, we referred to the isolation host information in GenBank whenever available and filtered out nonbacterial isolation hosts. We then standardized the host names following NCBI Taxonomy’s nomenclature [48]. Note that, while some phages are known to be polyvalent (multihost), only five phage entries were recorded with multiple host genera. Hence, for simplicity, we mapped each polyvalent phage to its host with the highest number of interacting phages in the dataset.

After preprocessing, host information was supplied for an additional 84 phages, thus totaling 15,823 phages across 279 host genera; the additional identified genus, *Silvanigrella*, was from the isolation host of MWH-Nonnen-W8red.

#### Identification of RBPs

Among the phage entries with host data, 15,158 entries have gene annotations, while the remaining 665 do not. For those with annotations, we selected the annotated RBPs using a regular expression (S1 Listing) and a manual exclusion list adapted from Boeckaerts *et al*. [49]; this exclusion list covers proteins related to RBPs but are not RBPs themselves (e.g., assembly and portal proteins). We also discarded sequences with undetermined amino acids (X).

Meanwhile, we ran the genomes of phages without annotation through Prokka [50], a wrapper tool for genome annotation. It first calls Prodigal [51] to predict the coding sequence coordinates. To identify the putative gene products, we configured Prokka [50] to refer to PHROG [52], a database of viral protein family clusters generated by employing hidden Markov model profile-profile comparisons for remote homology detection. PHROG [52] has also been used in previous studies that require the functional annotation of phage protein sequences [53, 54]. The annotated RBPs were selected following the same scheme described in the previous paragraph.

Afterwards, we discarded RBPs with lengths outside the interval [*Q*_1_ − · 1.5 *IQR, Q*_3_ + 1.5 *· IQR*], where *Q*_1_ is the first quartile, *Q*_3_ is the third quartile, and *IQR* is the interquartile range of the RBP lengths. This resulted in the removal of protein sequences longer than 1,587 amino acids. Figs 2A and 2B show the distribution of the lengths of the RBPs before and after this step, respectively.

**Fig 2.**
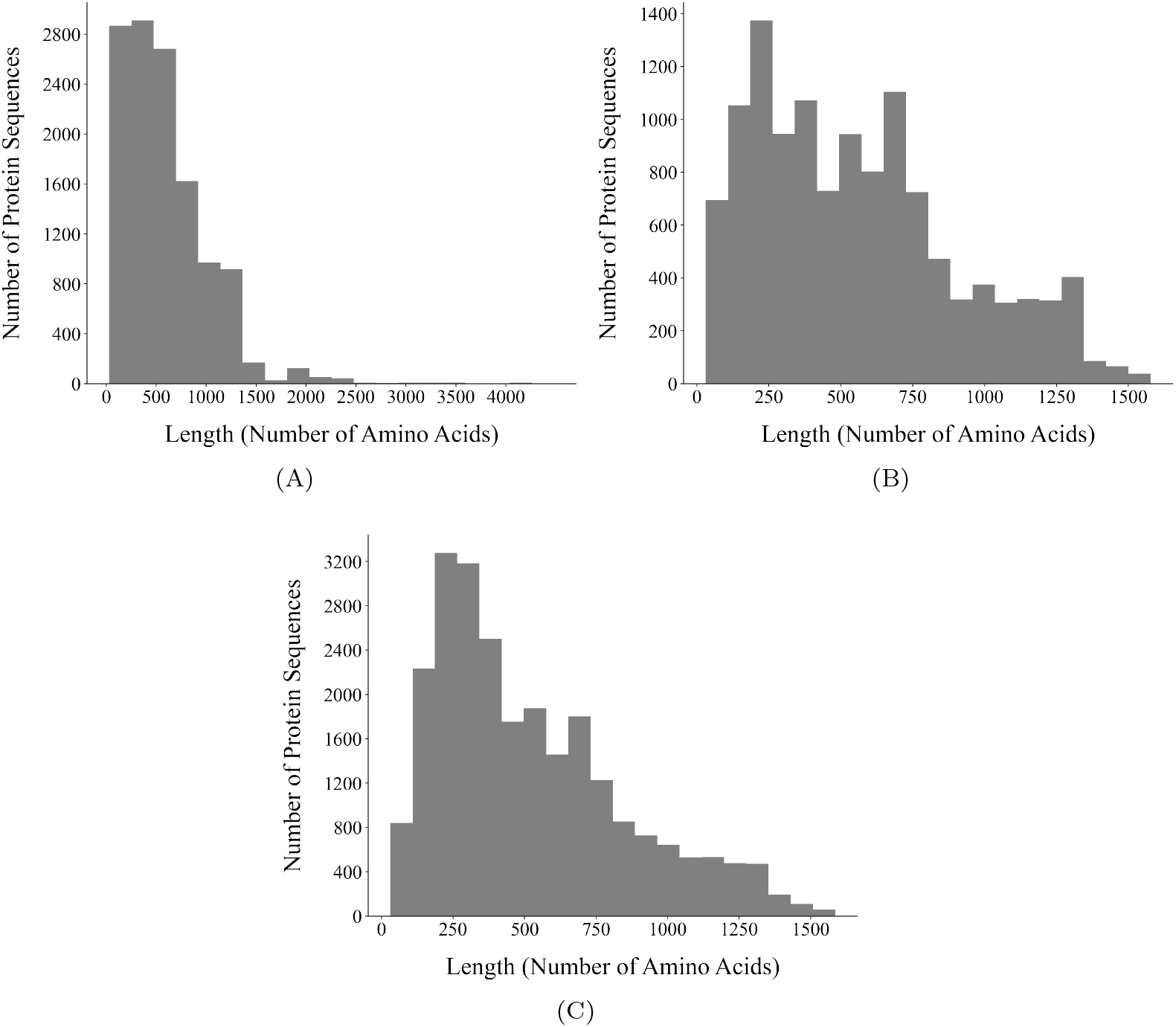
Distribution of the lengths of the RBPs. (A) Distribution of the lengths of the annotated RBPs selected based on annotation in GenBank and the functional annotation obtained using PHROG [52]. (B) Distribution of the lengths of the annotated RBPs after excluding those longer than 1,587 amino acids. This cutoff was set by defining outlying lengths as those outside the interval [*Q*_1_ − · 1.5 *IQR, Q*_3_ + 1.5 *· IQR*], where *Q*_1_ is the first quartile, *Q*_3_ is the third quartile, and *IQR* is the interquartile range of the RBP lengths. (C) Distribution of the lengths of all the RBPs in our dataset, including those computationally predicted via the approach proposed by Boeckaerts *et al*. [49].

Finally, to expand the list of RBPs in our dataset, we also considered proteins labeled as hypothetical by GenBank or uncharacterized by Prokka. Following Boeckaerts *et al*. [49], we encoded these sequences via the transformer-based protein language model ProtBert [31] and fed the generated embeddings to their proposed extreme gradient boosting model to computationally predict whether the hypothetical proteins are RBPs or not. In total, our dataset consists of 24,752 RBPs across 9,583 phages and 232 hosts (Table 1); the distribution of the lengths of these RBPs is presented in Fig 2C.

**Table 1.** Statistics on the identification of RBPs.

|  | Num. of RBPs | Num. of Phages | Num. of Hosts |
| --- | --- | --- | --- |
| GenBank | 11,133 | 5,805 | 167 |
| PHROG | 1,010 | 419 | 30 |
| Predicted | 12,609 | 5,941 | 192 |
| Total | 24,752 | 9,583 | 232 |

GenBank refers to the RBPs annotated in GenBank, PHROG refers to those selected based on the functional annotation obtained using PHROG [52]. *Predicted* refers to those computationally predicted via the approach proposed by Boeckeaerts *et al.* [49].

### Representation via protein language models

To generate dense vector representations (embeddings) of the RBP sequences, we explored seven pretrained protein language models as feature encoders: ESM [38], ESM-1b [38], ProtBert [31], ProtXLNet [31], ProtAlbert [31], and ProtT5 [31], and SeqVec [36]. Technical details about these protein language models are summarized in Table 2.

**Table 2.** Protein language models for generating receptor-binding protein embeddings.

| Model | Architecture | Pretraining Datasets | Vector length |
| --- | --- | --- | --- |
| SeqVec [36] | ELMo [37] | UniRef50 [40] | 1024 |
| ESM [38] | Transformer | UniParc [39] | 1280 |
| ESM-1b [38] | Transformer | UniRef50 [40] | 1280 |
| ProtBert [31] | Transformer | UniRef100 [40], BFD100 [41, 42] | 1024 |
| ProtXLNet [31] | Transformer | UniRef100 [40] | 1024 |
| ProtAlbert [31] | Transformer | UniRef100 [40] | 4096 |
| ProtT5 [31] | Transformer | UniRef50 [40], BFD100 [41, 42] | 1024 |

### Classifier building

We framed phage-host interaction prediction as a multiclass classification problem, with the protein embeddings of the RBPs as the input and the host (at the genus level) as the output. Including all 232 hosts in our dataset resulted in class imbalance, with a quarter of the hosts already accounting for 96.02% of the dataset entries. To mitigate this, we restricted the class labels to only the top 25% (i.e., 58) hosts associated with the most RBPs. The class labels are enumerated in S1 Table.

We divided our dataset into two sets *D*_1_ and *D*_2_, where *D*_1_ contains the RBPs with class labels belonging to the top 25% hosts and *D*_2_ contains the remaining entries. We then partitioned *D*_1_ following a stratified 70%-30% train-test split and appended *D*_2_ to the test set. As such, our test set includes RBPs with class labels outside the top 25% hosts, which we will refer to as the *others* class. This is to make the evaluation more reflective of real-world use cases, where we might encounter inputs not belonging to any class for which the model trained. In total, our training and test sets have 16,636 and 8,116 samples, respectively. Table 3 reports the training and test set statistics on the top 10 hosts associated with the most RBPs; the complete statistics are given in S1 Table.

**Table 3.**
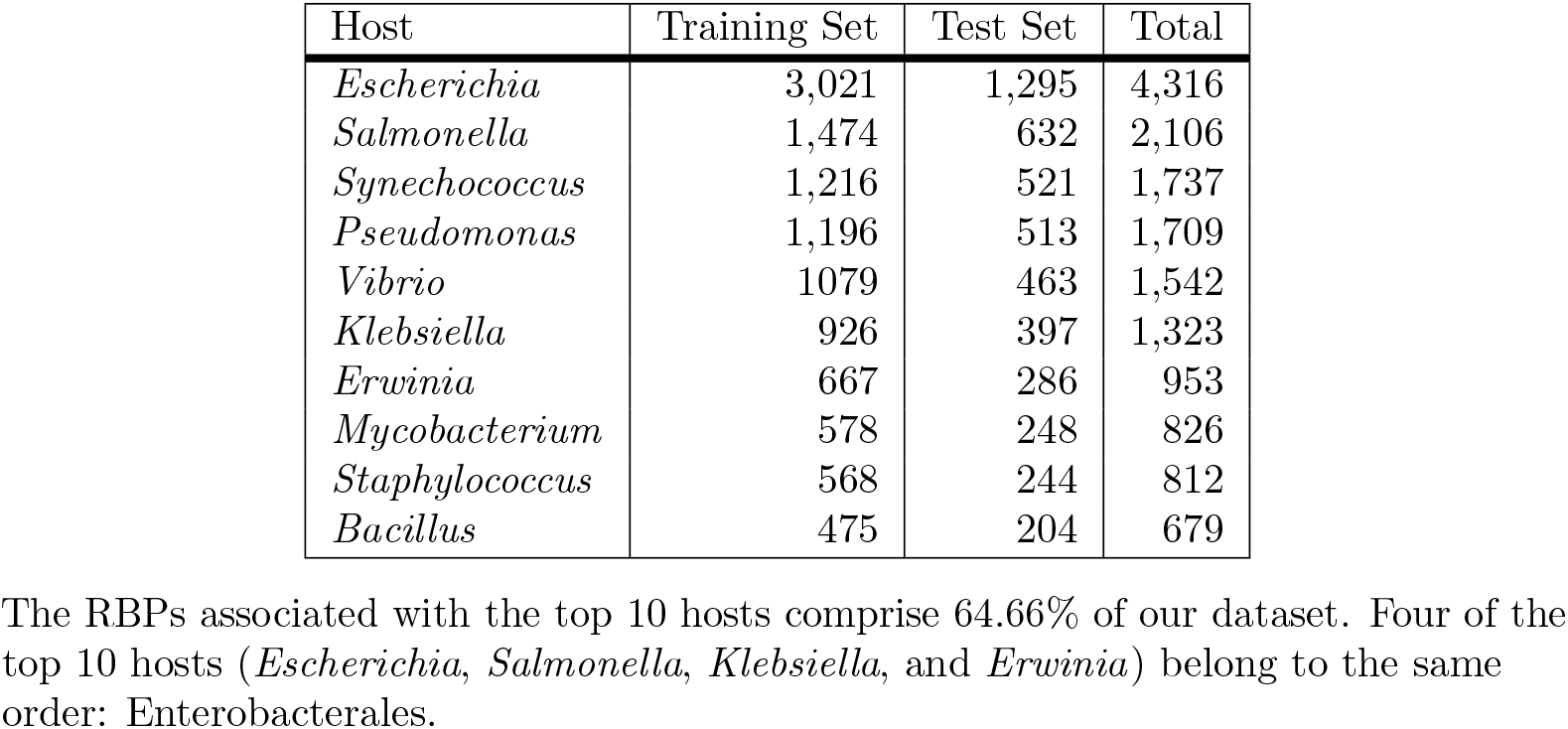
Number of training and test samples for the top 10 hosts associated with the most receptor-binding proteins (RBPs).

Afterwards, we fed the protein embeddings to a random forest classifier. In a further attempt to address class imbalance, we employed a weighted random forest model that imposes higher misclassification penalties for minority classes [55, 56]. Formally, let *N* be the total number of samples in the training set, *c* be the number of classes, and *n* be the number of training samples under a given class. The misclassification penalty *w* for this class is given by Eq (1).

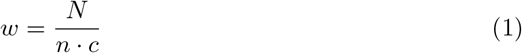

To optimize the weighted F1 score, we conducted hyperparameter tuning with five-fold stratified cross-validation. The hyperparameter space is as follows (the optimal values are in bold): number of trees (50, 100, **150**, 200), number of features to consider in determining the best split (log_2_, **square root**), minimum number of samples to split an internal node (**2**, 3, 4), and minimum number of samples to be a leaf node (**1**, 2, 3, 4).

### Performance evaluation

In order to factor in our model’s confidence in its prediction, we introduced a confidence threshold *k*. Let *p*_1_ and *p*_2_ be the highest and second-highest predicted class probabilities, respectively, for an input RBP. This input is classified under its predicted class label if and only if *p*_1_ *− p*_2_ ≥ *k*. If *p*_1_ *− p*_2_ *< k*, then it is classified as *others* since the model is ambiguous about its classification. S2 Table defines the true and false positive and negative outcomes in view of this scheme.

We evaluated our model’s performance using weighted precision, recall, and F1, with the weights corresponding to the class sizes. Formally, let *N* be the total number of samples in the test set, *C* be the set of class labels including the *others* class, *n*_*c*_ be the number of test samples under class *c*. Let *TP*_*c*_, *TN*_*c*_, *FP*_*c*_, and *FN*_*c*_ be the number of true positive, true negative, false positive, and false negative outcomes for class *c*. The definitions of the evaluation metrics are given in Eqs (2) to (4).

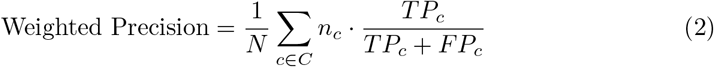

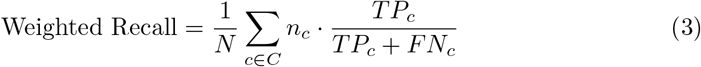

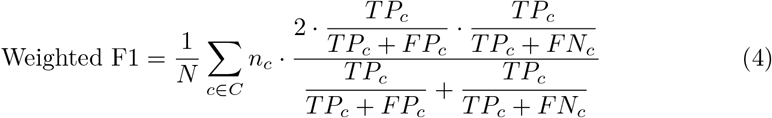

## Results

### Transformer-based embeddings of RBPs outperformed handcrafted genomic and protein features for phage-host interaction prediction

We compared the performance of our model with that of the state-of-the-art phage-host interaction prediction tool by Boeckaerts *et al*. [15]; its feature set consists of 218 handcrafted features, 133 of which are derived from the genomic sequences and 85 from the RBP sequences. We evaluated their performance across different confidence thresholds ranging from *k* = 60% to 100% in steps of 10%.

While representing RBPs via protein embeddings and via these handcrafted features yielded similar weighted precision scores (S3 Table), the use of protein embeddings improved the weighted F1 and recall across all tested confidence thresholds (Table 4 and S4 Table). Among the evaluated protein language models, the highest performance was obtained with the autoencoder model ProtT5, followed by ESM-1b and ESM. In particular, utilizing the ProtT5 embeddings outperformed the handcrafted features by around 3% to 4% in terms of weighted F1 and recall; the per-class evaluation results are reported in S5 Table to S9 Table. Meanwhile, the smallest performance increase was registered by the autoregressive model ProtXLNet.

### Integrating handcrafted features did not significantly increase performance

To determine the extent to which the further integration of handcrafted sequence properties can potentially improve the predictive power of our best-performing model, we combined the ProtT5 embeddings of the RBPs with the sequence properties that registered the highest Gini importance after training the model by Boeckaerts *et al*. [15] on our dataset. We also examined the performance if the sequence properties were limited to those extracted from protein sequences.

**Table 4.** Model performance in terms of weighted F1.

|  | 60% | 70% | 80% | 90% | 100% |
| --- | --- | --- | --- | --- | --- |
| Boeckaerts *et al.* [15] | 59.35% | 53.41% | 47.20% | 38.97% | 21.61% |
| SeqVec | 60.64% | 54.90% | 49.12% | 40.52% | 23.20% |
| ESM | 62.21% | 57.13% | 51.18% | 42.70% | **<u>25.18%</u>** |
| ESM-1b | 62.27% | 57.16% | 51.29% | 42.66% | 25.04% |
| ProtBert | 60.58% | 55.27% | 49.29% | 41.07% | 24.59% |
| ProtXLNet | 60.18% | 54.54% | 48.46% | 40.20% | 23.92% |
| ProtAlbert | 60.45% | 54.82% | 48.51% | 40.57% | 23.80% |
| ProtT5 | **<u>62.95%</u>** | **<u>57.51%</u>** | **<u>51.98%</u>** | **<u>43.05%</u>** | 24.82% |

The header row refers to the confidence thresholds at which we evaluated model performance; these confidence thresholds range from $k = 60\%$ to $100\%$ in steps of $10\%$ . The highest weighted F1 scores are given in bold and underlined.

As reported in Tables 5 to 8, the increase in the weighted F1 scores after integrating these handcrafted sequence properties was consistently below 1.33% across all tested confidence thresholds. Furthermore, the increase in precision and recall was also consistently below 1.40% (S10 Table to S13 Table) and 1.41% (S14 Table to S17 Table), respectively. These results suggest that the ProtT5 embeddings may already be capturing these sequence properties.

**Table 5.**
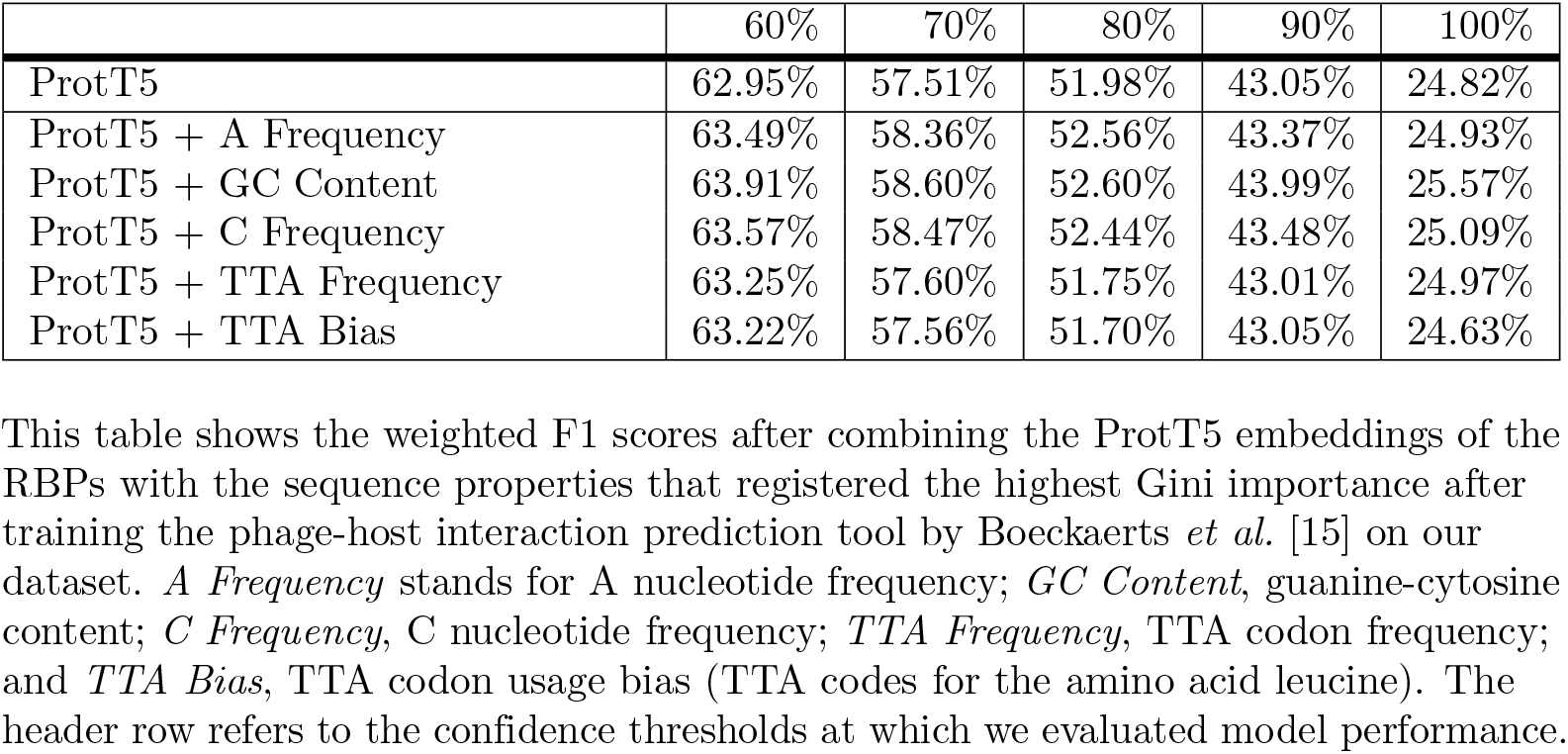
Weighted F1 scores after integrating handcrafted sequence properties to the vector representations of the RBPs.

|  | 60% | 70% | 80% | 90% | 100% |
| --- | --- | --- | --- | --- | --- |
| ProtT5 | 62.95% | 57.51% | 51.98% | 43.05% | 24.82% |
| ProtT5 + A Frequency | 63.49% | 58.36% | 52.56% | 43.37% | 24.93% |
| ProtT5 + GC Content | 63.91% | 58.60% | 52.60% | 43.99% | 25.57% |
| ProtT5 + C Frequency | 63.57% | 58.47% | 52.44% | 43.48% | 25.09% |
| ProtT5 + TTA Frequency | 63.25% | 57.60% | 51.75% | 43.01% | 24.97% |
| ProtT5 + TTA Bias | 63.22% | 57.56% | 51.70% | 43.05% | 24.63% |

This table shows the weighted F1 scores after combining the ProtT5 embeddings of the RBPs with the sequence properties that registered the highest Gini importance after training the phage-host interaction prediction tool by Boeckaerts *et al.* [15] on our dataset. *A Frequency* stands for A nucleotide frequency; *GC Content*, guanine-cytosine content; *C Frequency*, C nucleotide frequency; *TTA Frequency*, TTA codon frequency; and *TTA Bias*, TTA codon usage bias (TTA codes for the amino acid leucine). The header row refers to the confidence thresholds at which we evaluated model performance.

## Discussion

### Representing receptor-binding proteins RBPs using protein language models

The main novelty of our study lies in our use of protein language models to obtain dense vector encodings of receptor-binding proteins, which are known to be major determinants of phage-host specificity [26–29].

**Table 6.** Weighted F1 scores after integrating handcrafted protein sequence properties to the vector representations of the receptor-binding proteins (RBPs).

|  | 60% | 70% | 80% | 90% | 100% |
| --- | --- | --- | --- | --- | --- |
| ProtT5 | 62.95% | 57.51% | 51.98% | 43.05% | 24.82% |
| ProtT5 + K Frequency | 63.00% | 57.49% | 52.00% | 43.13% | 25.07% |
| ProtT5 + pI | 62.90% | 57.36% | 51.76% | 43.01% | 25.27% |
| ProtT5 + Z4 | 62.86% | 57.40% | 51.48% | 43.06% | 24.97% |
| ProtT5 + % of Exposed SA | 62.78% | 57.65% | 51.64% | 42.91% | 25.42% |
| ProtT5 + Mol. Weight | 62.98% | 57.74% | 51.79% | 42.87% | 25.15% |

This table shows the weighted F1 scores after combining the ProtT5 embeddings of the RBPs with the protein sequence properties that registered the highest Gini importance after training the phage-host interaction prediction tool by Boeckaerts *et al.* [15] on our dataset. *K Frequency* stands for lysine frequency; *pI*, isoelectric point; *Z4*, fourth protein Z-scale [57], which is related to the heat of formation, hardness, electronegativity, and electrophilicity; *% of Exposed SA*, percentage of residues with exposed solvent accessibility; and *Mol. Weight*, molecular weight. The header row refers to the confidence thresholds at which we evaluated model performance.

**Table 7.** Weighted F1 scores after integrating the top *n* handcrafted sequence properties to the vector representations of the receptor-binding proteins (RBPs).

|  | 60% | 70% | 80% | 90% | 100% |
| --- | --- | --- | --- | --- | --- |
| ProtT5 | 62.95% | 57.51% | 51.98% | 43.05% | 24.82% |
| ProtT5 + Top 1 | 63.49% | 58.36% | 52.56% | 43.37% | 24.93% |
| ProtT5 + Top 2 | 64.26% | 58.76% | 52.70% | 43.75% | 25.13% |
| ProtT5 + Top 3 | 64.17% | 58.83% | 52.81% | 44.20% | 25.33% |
| ProtT5 + Top 4 | 64.03% | 58.83% | 52.67% | 43.85% | 24.44% |
| ProtT5 + Top 5 | 64.01% | 58.68% | 52.77% | 44.21% | 24.80% |

*ProtT5 + Top $n$* (where $n = 1$ to 5) indicates that the ProtT5 embeddings of the RBPs were combined with the $n$ sequence properties that registered the highest Gini importance after training the phage-host interaction prediction tool by Boeckaerts *et al.* [15] on our dataset. These protein sequence properties (in order of decreasing importance) are A nucleotide frequency, guanine-cytosine content, C nucleotide frequency, TTA codon frequency, and TTA codon usage bias (TTA codes for the amino acid leucine). The header row refers to the confidence thresholds at which we evaluated model performance.

Vectorizing sequences by representation learning requires only the sequences themselves, discarding the need to derive additional alignment or structural information and eliminating the difficulty of selecting from a wide array of potentially informative signals [30]. Aside from these general advantages of representation learning over manual feature engineering, our experiments also showed that the use of protein embeddings improves phage-host interaction prediction and outperforms handcrafted genomic and protein features. In particular, utilizing the transformer-based autoencoder model ProtT5 resulted in the best performance, increasing the weighted F1 scores by 3% to 4% across all tested confidence thresholds.

**Table 8.** Weighted F1 scores after integrating the top *n* handcrafted protein sequence properties to the vector representations of the receptor-binding proteins (RBPs).

|  | 60% | 70% | 80% | 90% | 100% |
| --- | --- | --- | --- | --- | --- |
| ProtT5 | 62.95% | 57.51% | 51.98% | 43.05% | 24.82% |
| ProtT5 + Protein Top 1 | 63.00% | 57.49% | 52.00% | 43.13% | 25.07% |
| ProtT5 + Protein Top 2 | 62.96% | 57.57% | 51.79% | 42.87% | 25.17% |
| ProtT5 + Protein Top 3 | 62.95% | 57.47% | 51.38% | 43.09% | 25.19% |
| ProtT5 + Protein Top 4 | 62.92% | 57.63% | 51.67% | 43.36% | 25.34% |
| ProtT5 + Protein Top 5 | 62.79% | 57.82% | 51.83% | 42.86% | 24.80% |

*ProtT5 + Protein Top $n$* (where $n = 1$ to 5) indicates that the ProtT5 embeddings of the RBPs were combined with the $n$ protein sequence properties that registered the highest Gini importance after training the phage-host interaction prediction tool by Boeckaerts *et al.* [15] on our dataset. These sequence properties (in order of decreasing importance) are lysine frequency, isoelectric point, fourth protein Z-scale [57] (which is related to the heat of formation, hardness, electronegativity, and electrophilicity), percentage of residues with exposed solvent accessibility, and molecular weight. The header row refers to the confidence thresholds at which we evaluated model performance.

### Our embeddings-based model captures a complex combination of features

While protein embeddings have been shown to improve performance in several prototypical bioinformatics tasks [34, 58, 59], their interpretability remains a challenge [60]. The most common approach is to project the embeddings onto a low-dimensional space via nonlinear dimensionality reduction techniques [30, 31, 36, 61], such as *t*-distributed stochastic neighbor embedding (*t*-SNE) [62] and uniform manifold approximation and projection (UMAP) [63], and check for the presence of formed clusters. Vig *et al*. [64] analyzed the internal model representations of ProtBert, ProtAlbert, and ProtXLNet to demonstrate how attention learns protein folding structure and targets binding sites.

These approaches have been employed to establish that protein embeddings carry information on salient physicochemical, structural, and functional properties of protein sequences [33]. For instance, SeqVec and the ProtTrans and ESM families of language models, which we explored in our work, have been shown to capture properties such as hydrophobicity, charge, polarity, and molecular weight [31, 36, 38]. In our experiments, we found no significant performance improvement after combining the ProtT5 embeddings with selected handcrafted features, suggesting that the embeddings may already be capturing guanine-cytosine content [15, 65–67], codon usage bias [15, 68–70], and other important signals of phage-host interaction that emerge from the close coexistence and coevolution of phages and their bacterial hosts.

To visualize the geometry of the embeddings and attempt to identify the biophysical features that they capture in the context of phage-host interaction prediction, we represented each RBP as a vector whose components are the *ℓ* components of its ProtT5 embedding with the highest Gini importance after training our model. We then employed *t*-SNE and UMAP [63] to project the set of these vectors onto a two-dimensional space. We colored the points based on the sequence properties with the highest Gini importance after training the model by Boeckaerts *et al*. [15] on our dataset. Figures 3 and 4 show the resulting plots when *ℓ* is set to 100.

**Fig 3.**
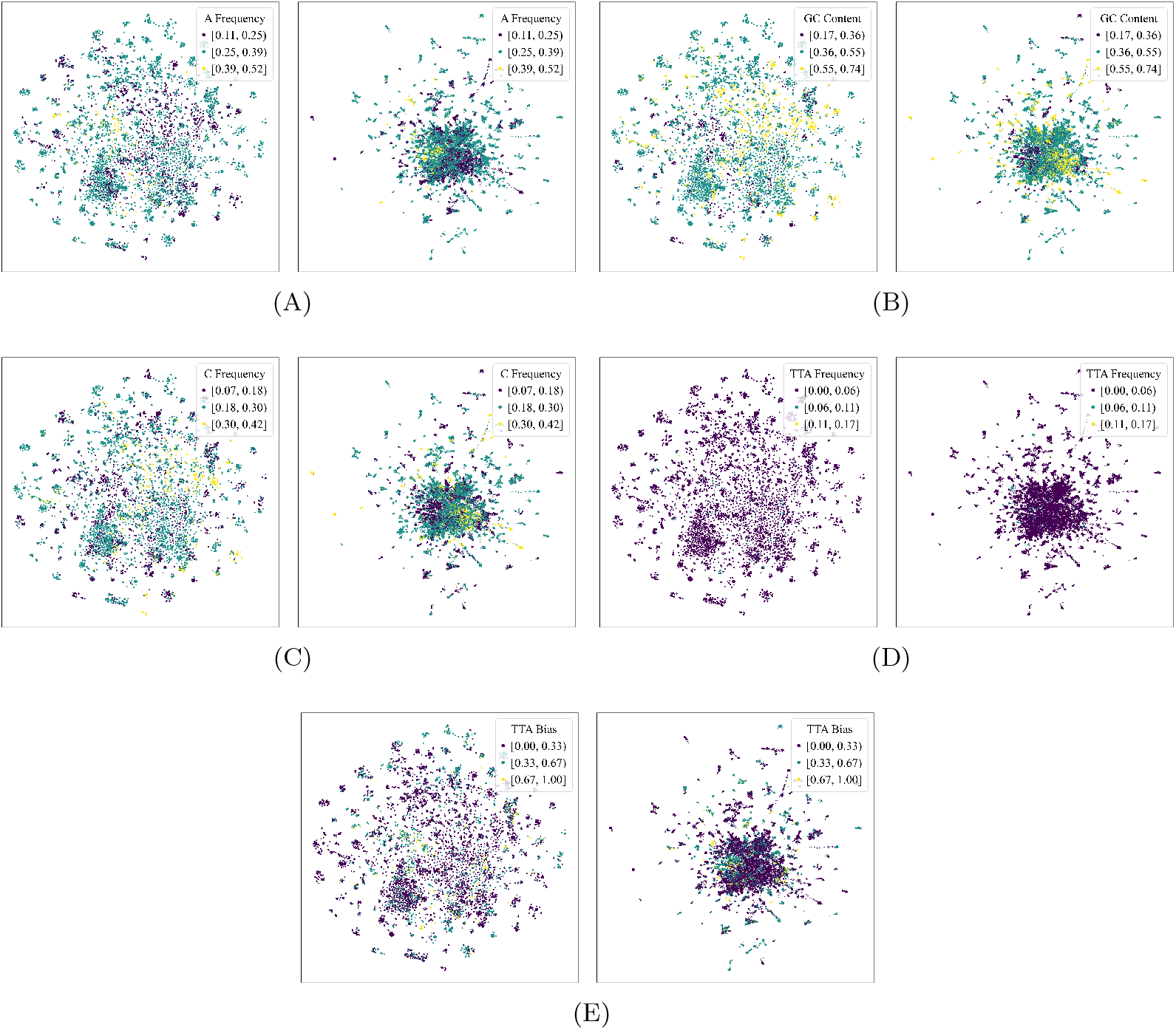
*t* -distributed stochastic neighbor embedding (*t* -SNE) and uniform manifold approximation and projection (UMAP) plots of the ProtT5 embeddings, colored based on handcrafted sequence properties. The *t* -SNE projections (left of each subfigure) were generated by setting the perplexity to 50, the number of iterations to 1000, the initialization to principal component analysis, and the learning rate to the number of samples divided by 12, following Kobak and Berens [71]. The UMAP projections (right of each subfigure) were generated by setting the number of neighbors to 100 and the minimum distance between embedded points to 0.7. Each point corresponds to the two-dimensional projection of a subvector of a receptor-binding protein’s ProtT5 embedding, the components of which are the *ℓ* components with the highest Gini importance after training our phage-host interaction prediction model (in this figure, *ℓ* = 100). The points were colored based on the sequence properties with the highest Gini importance after training the model by Boeckaerts *et al*. [15] on our dataset; these properties are as follows: (A) A nucleotide frequency, (B) guanine-cytosine content, (C) C nucleotide frequency, (D) TTA codon frequency, and (E) TTA codon usage bias (TTA codes for the amino acid leucine).

**Fig 4.**
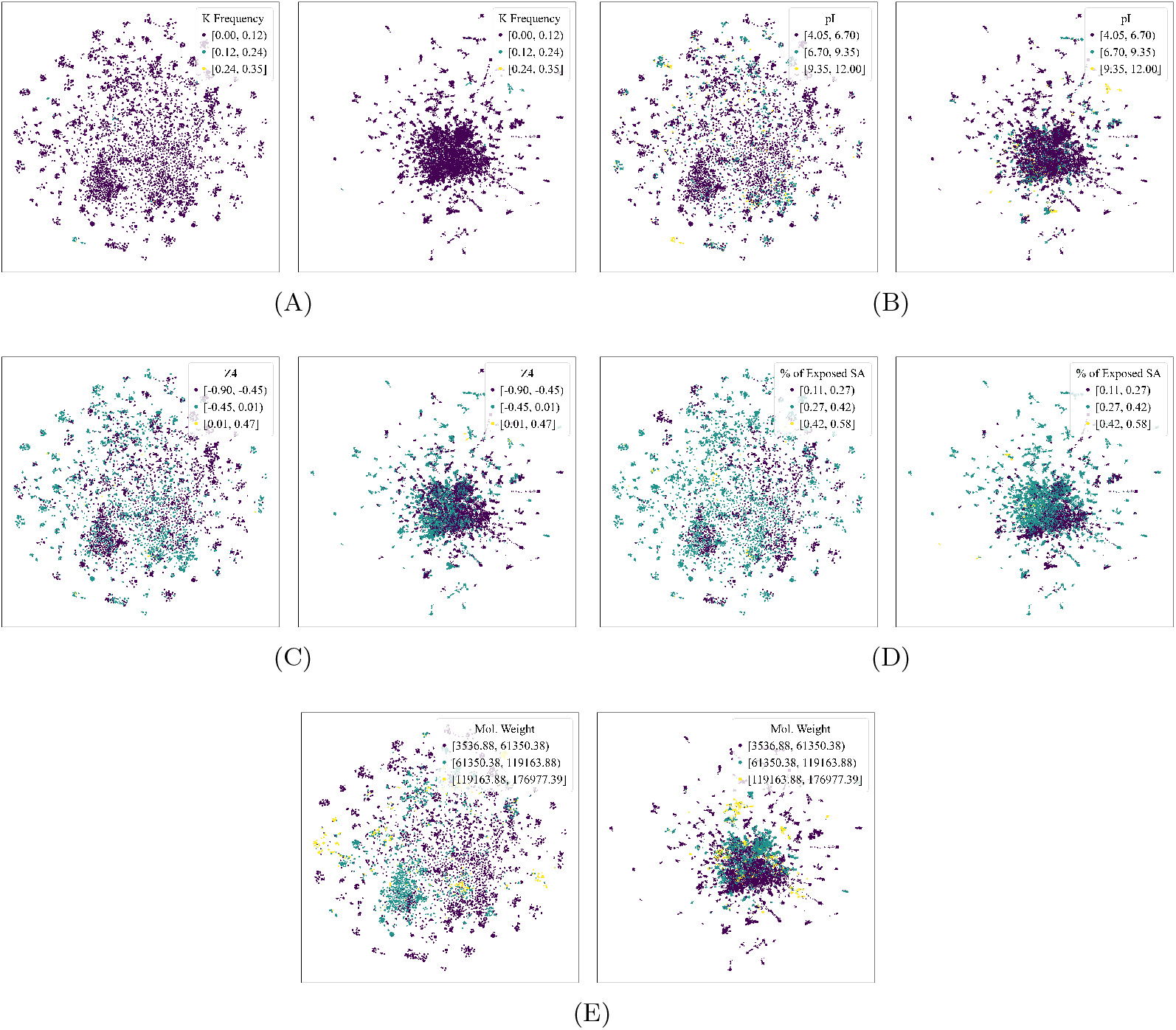
*t* -distributed stochastic neighbor embedding (*t* -SNE) and uniform manifold approximation and projection (UMAP) plots of the ProtT5 embeddings, colored based on handcrafted protein sequence properties. The *t* -SNE projections (left of each subfigure) were generated by setting the perplexity to 50, the number of iterations to 1000, the initialization to principal component analysis, and the learning rate to the number of samples divided by 12, following Kobak and Berens [71]. The UMAP projections (right of each subfigure) were generated by setting the number of neighbors to 100 and the minimum distance between embedded points to 0.7. Each point corresponds to the two-dimensional projection of a subvector of a receptor-binding protein’s ProtT5 embedding, the components of which are the *ℓ* components with the highest Gini importance after training our phage-host interaction prediction model (in this figure, *ℓ* = 100). The points were colored based on the protein sequence properties with the highest Gini importance after training the model by Boeckaerts *et al*. [15] on our dataset; these properties are as follows: (A) lysine frequency, (B) isoelectric point, (C) fourth protein Z-scale [57] (which is related to the heat of formation, hardness, electronegativity, and electrophilicity), (D) percentage of residues with exposed solvent accessibility, and (E) molecular weight.

Experimenting with different values of *ℓ*, we observed that, although there were instances wherein local clusters appeared to form, the points generally did not show clear separation boundaries, suggesting that the components of the embeddings do not squarely correspond to the individual sequence features under consideration. We hypothesize that, while embeddings certainly carry biophysical markers of phage-host interaction, their components cannot necessarily be disentangled and mapped to specific features. Instead, they likely capture these features in some complex combination. Investigating the exact fashion in which these features are combined is a possible research direction for improving interpretability.

## Conclusion

In this study, we capitalized on representation learning to automatically encode receptor-binding protein sequences into meaningful dense embeddings. To this end, we extensively tested different protein language models and built a random forest model for phage-host interaction prediction. Our experiments showed that the use of embeddings of receptor-binding proteins presents improvements over handcrafted genomic and protein sequence features, with the highest performance obtained using the transformer-based autoencoder model ProtT5. Moreover, these protein embeddings are able to capture complex combinations of biological features given only the raw sequences, without the need to supply additional alignment or structural information.

Our work makes the simplifying assumption that all the RBPs of a given phage are specific to one host. Albeit significantly less common than single-host phages, some phages are known to possess multiple RBPs, with the RBPs possibly adsorbing with different bacteria. For example, the polyvalent bacteriophage ΦK64-1 has eleven known RBPs targeting a wide spectrum of *Klebsiella* capsular types [72]. However, to the best of our knowledge, there are currently no existing datasets that map individual RBPs to their target hosts.

The receptor-binding proteins considered in our study are also limited to those of tailed phages belonging to the order *Caudovirales*, which constitute around 96% of all known phages [73]. It may also be interesting to explore a similar approach for the computational prediction of interaction between non-tailed phages and their hosts.

Further future directions include improving the interpretability of protein embeddings and incorporating other mechanisms related to phage-host interaction (e.g., restriction-modification and CRISPR-Cas systems), as well as host sequence information possibly encoded as dense embeddings.

## Data availability statement

The data and source code for the experiments and analysis are available at https://github.com/bioinfodlsu/phage-host-prediction.

## Supporting information

**S1 Listing. Regular expression for the selection of annotated receptor-binding proteins**. We modified the regular expression from Boeckaerts *et al*. [49] to better accommodate typographical errors and naming variations in the gene product annotations in GenBank.

~~~
tail?(.?|\s*)(?:spike?|fib(?:er|re))|recept(?:o|e)r(.?|\s*)(?:bind|recogn) .*(?:protein)?|(?<!\w)RBP(?!a)
~~~

**S1 Table.**
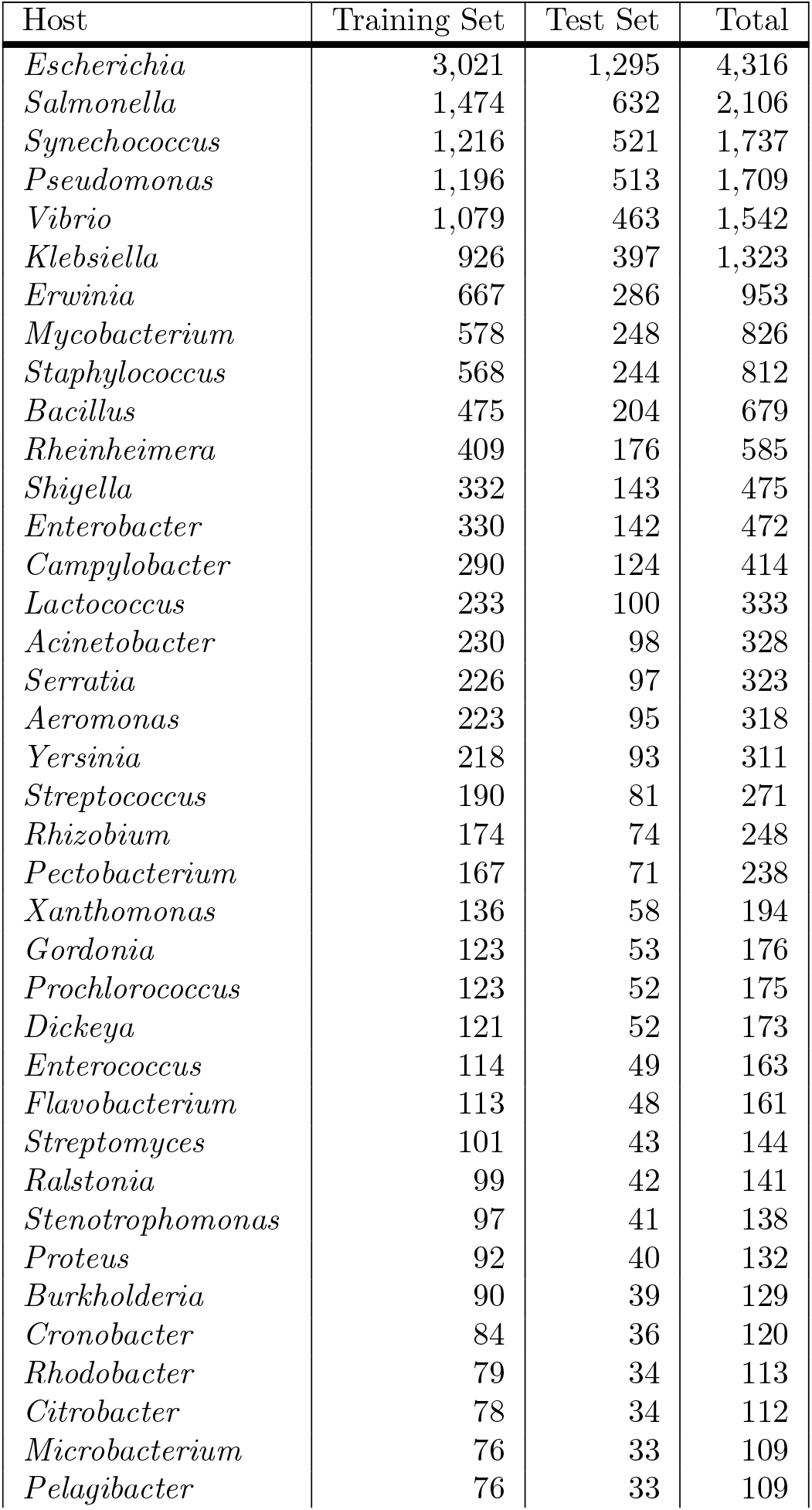
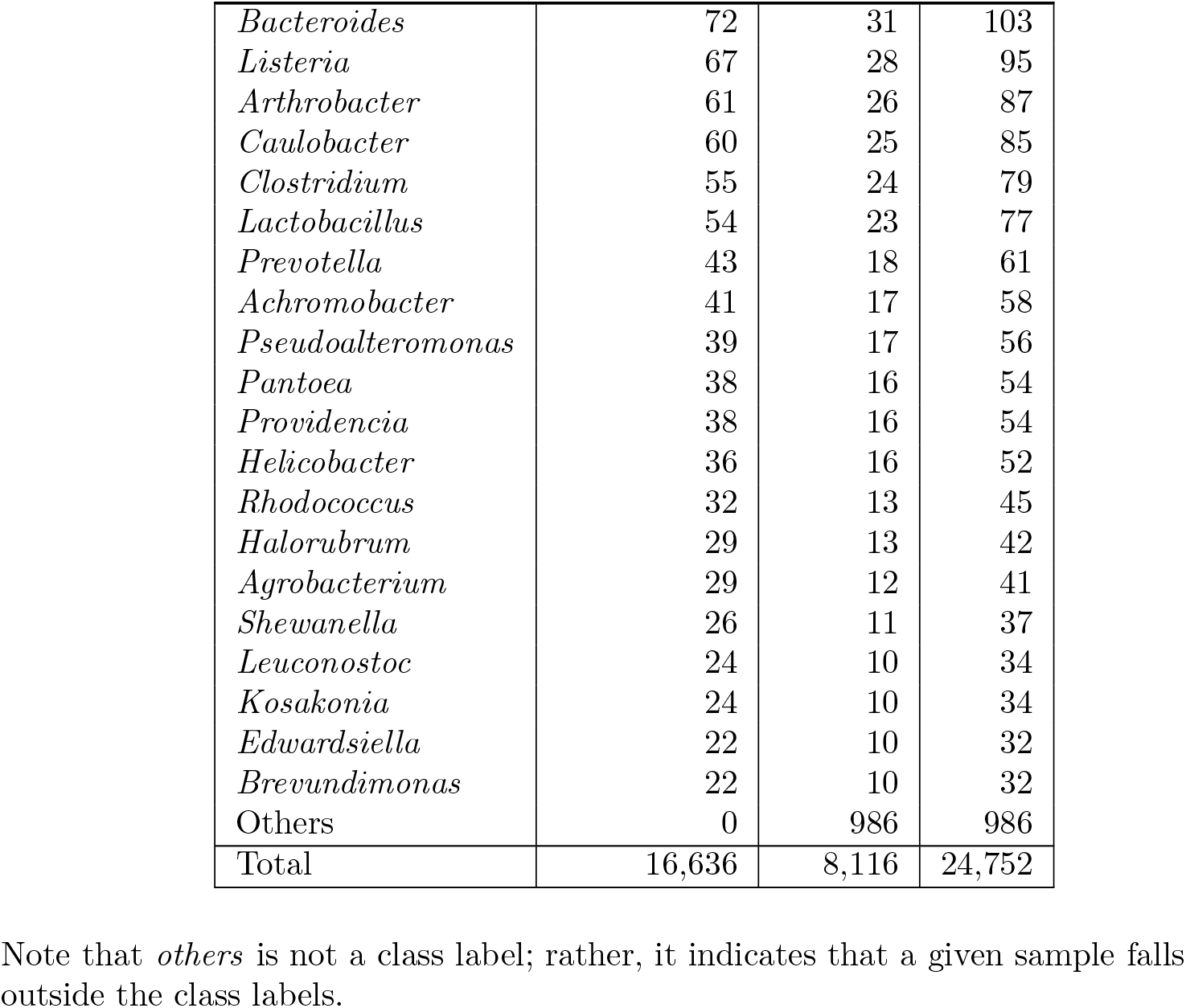
Number of training and test samples for all class labels.

**S2 Table.** Definition of true and false positive and negative outcomes.

| $p_1 - p_2$ | True Label | Predicted Label (Original) | Predicted Label (In View of $p_1 - p_2$) | Evaluation Outcome |
| --- | --- | --- | --- | --- |
| $\geq k$ | *X* | *X* | *X* | TP for *X* |
|  | *Y* | *X* | *X* | FN for *Y*<br>FP for *X* |
|  | *others* | *X* | *X* | FN for *others*<br>FP for *X* |
| $< k$ | *X* | *X* | *others* | FN for *X*<br>FP for *others* |
|  | *Y* | *X* | *others* | FN for *Y*<br>FP for *others*<br>TN for *X* |
|  | *others* | *X* | *others* | TP for *others*<br>TN for *X* |

In the first column, $p_1$ and $p_2$ are the highest and second highest class probabilities for a given sample, and $k$ is a confidence threshold. A sample is classified under its (original) predicted class label if and only if $p_1 - p_2 \geq k$ . If $p_1 - p_2 < k$ , then it is classified as *others*. In the second to last columns, *X* and *Y* refer to two distinct class labels; *others* indicates that the sample falls outside the class labels. In the last column, *TP*, *FP*, *TN*, and *FN* stand for *true positive*, *false positive*, *true negative*, and *false negative*, respectively.

**S3 Table.** Model performance in terms of weighted precision.

|  | 60% | 70% | 80% | 90% | 100% |
| --- | --- | --- | --- | --- | --- |
| Boeckaerts *et al.* [15] | 85.55% | 84.88% | 84.05% | 83.58% | 73.09% |
| SeqVec | 85.65% | 84.56% | 84.26% | **83.83%** | 73.20% |
| ESM | 85.31% | 84.62% | 83.66% | 82.68% | 76.51% |
| ESM-1b | 84.71% | **84.99%** | **84.40%** | 83.42% | 76.89% |
| ProtBert | 84.96% | 84.55% | 84.20% | 83.69% | 73.74% |
| ProtXLNet | **85.66%** | 84.67% | 84.06% | 82.80% | 77.07% |
| ProtAlbert | 84.91% | 84.61% | 83.97% | 83.03% | 76.06% |
| ProtT5 | 85.43% | 84.98% | 84.32% | 83.51% | **77.23%** |

The header row refers to the confidence thresholds at which we evaluated model performance. These confidence thresholds range from $k = 60\%$ to $100\%$ in steps of $10\%$ . The highest weighted precision scores are given in bold and underlined.

**S4 Table.** Model performance in terms of weighted recall.

|  | 60% | 70% | 80% | 90% | 100% |
| --- | --- | --- | --- | --- | --- |
| Boeckaerts *et al.* [15] | 55.48% | 49.84% | 44.50% | 37.81% | 24.84% |
| SeqVec | 56.73% | 51.22% | 45.97% | 38.87% | 25.89% |
| ESM | 58.33% | 53.38% | 47.92% | 40.69% | **27.34%** |
| ESM-1b | 58.49% | 53.30% | 47.86% | 40.59% | 27.24% |
| ProtBert | 56.65% | 51.60% | 46.16% | 39.48% | 27.02% |
| ProtXLNet | 56.32% | 50.95% | 45.50% | 38.50% | 26.56% |
| ProtAlbert | 56.67% | 51.21% | 45.53% | 38.90% | 26.38% |
| ProtT5 | **59.15%** | **53.72%** | **48.57%** | **41.03%** | 27.16% |

The header row refers to the confidence thresholds at which we evaluated model performance. These confidence thresholds range from $k = 60\%$ to $100\%$ in steps of $10\%$ . The highest weighted recall scores are given in bold and underlined.

**S5 Table.**
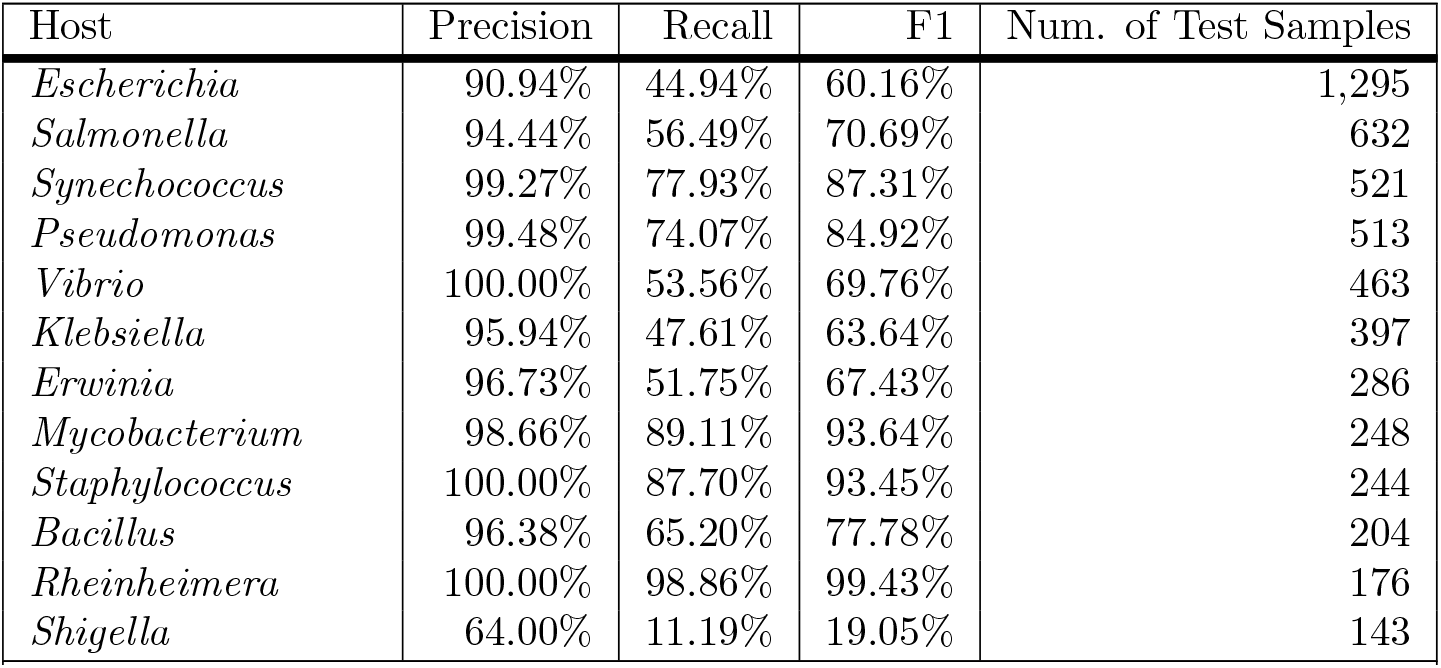
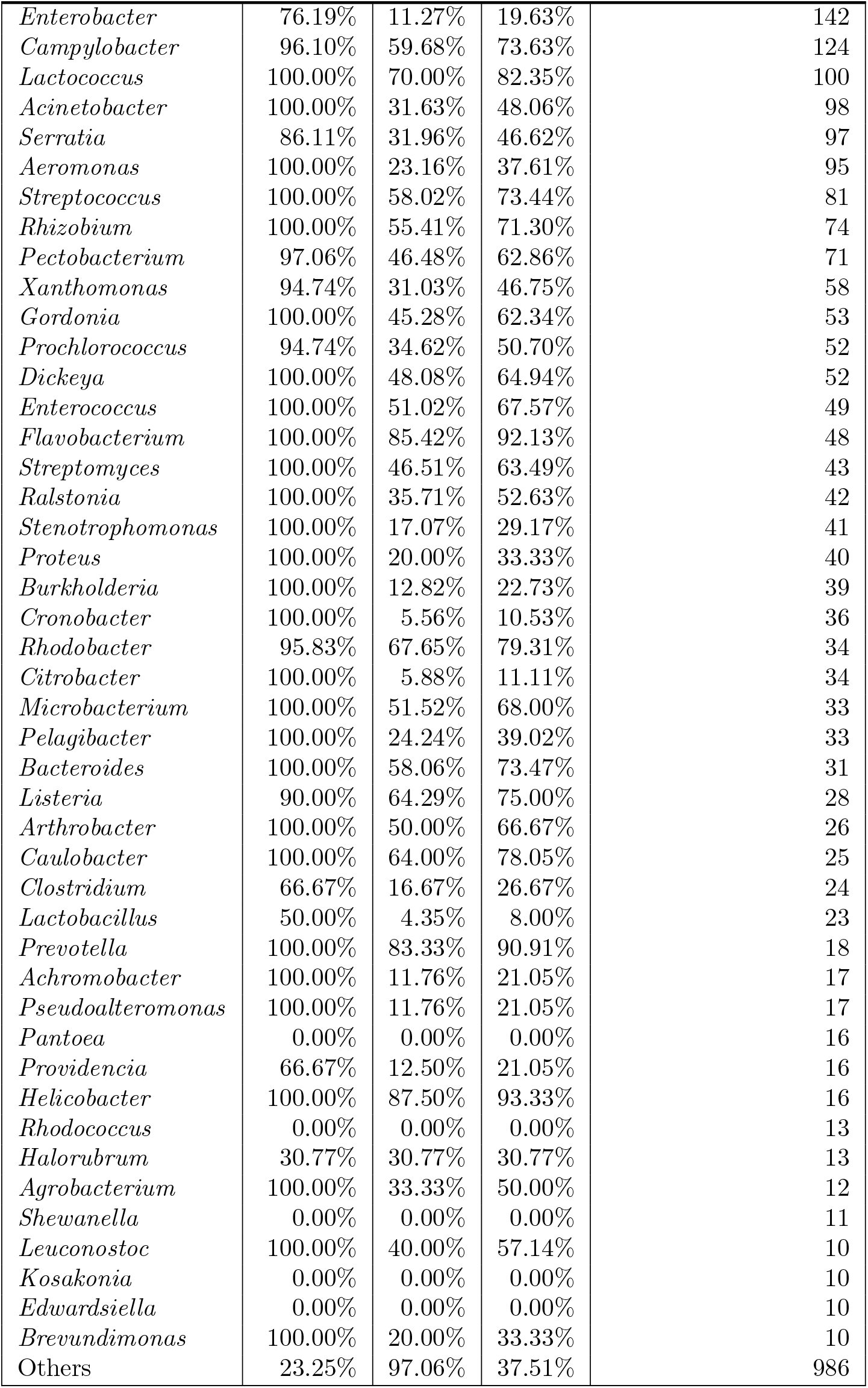
Per-class evaluation results of using ProtT5 embeddings at confidence threshold *k* = 60%.

| Host | Precision | Recall | F1 | Num. of Test Samples |
| --- | --- | --- | --- | --- |
| *Escherichia* | 90.94% | 44.94% | 60.16% | 1,295 |
| *Salmonella* | 94.44% | 56.49% | 70.69% | 632 |
| *Synechococcus* | 99.27% | 77.93% | 87.31% | 521 |
| *Pseudomonas* | 99.48% | 74.07% | 84.92% | 513 |
| *Vibrio* | 100.00% | 53.56% | 69.76% | 463 |
| *Klebsiella* | 95.94% | 47.61% | 63.64% | 397 |
| *Erwinia* | 96.73% | 51.75% | 67.43% | 286 |
| *Mycobacterium* | 98.66% | 89.11% | 93.64% | 248 |
| *Staphylococcus* | 100.00% | 87.70% | 93.45% | 244 |
| *Bacillus* | 96.38% | 65.20% | 77.78% | 204 |
| *Rheinheimera* | 100.00% | 98.86% | 99.43% | 176 |
| *Shigella* | 64.00% | 11.19% | 19.05% | 143 |
| Continued on the next page |  |  |  |  |
| *Enterobacter* | 76.19% | 11.27% | 19.63% | 142 |
| *Campylobacter* | 96.10% | 59.68% | 73.63% | 124 |
| *Lactococcus* | 100.00% | 70.00% | 82.35% | 100 |
| *Acinetobacter* | 100.00% | 31.63% | 48.06% | 98 |
| *Serratia* | 86.11% | 31.96% | 46.62% | 97 |
| *Aeromonas* | 100.00% | 23.16% | 37.61% | 95 |
| *Streptococcus* | 100.00% | 58.02% | 73.44% | 81 |
| *Rhizobium* | 100.00% | 55.41% | 71.30% | 74 |
| *Pectobacterium* | 97.06% | 46.48% | 62.86% | 71 |
| *Xanthomonas* | 94.74% | 31.03% | 46.75% | 58 |
| *Gordonia* | 100.00% | 45.28% | 62.34% | 53 |
| *Prochlorococcus* | 94.74% | 34.62% | 50.70% | 52 |
| *Dickeya* | 100.00% | 48.08% | 64.94% | 52 |
| *Enterococcus* | 100.00% | 51.02% | 67.57% | 49 |
| *Flavobacterium* | 100.00% | 85.42% | 92.13% | 48 |
| *Streptomyces* | 100.00% | 46.51% | 63.49% | 43 |
| *Ralstonia* | 100.00% | 35.71% | 52.63% | 42 |
| *Stenotrophomonas* | 100.00% | 17.07% | 29.17% | 41 |
| *Proteus* | 100.00% | 20.00% | 33.33% | 40 |
| *Burkholderia* | 100.00% | 12.82% | 22.73% | 39 |
| *Cronobacter* | 100.00% | 5.56% | 10.53% | 36 |
| *Rhodobacter* | 95.83% | 67.65% | 79.31% | 34 |
| *Citrobacter* | 100.00% | 5.88% | 11.11% | 34 |
| *Microbacterium* | 100.00% | 51.52% | 68.00% | 33 |
| *Pelagibacter* | 100.00% | 24.24% | 39.02% | 33 |
| *Bacteroides* | 100.00% | 58.06% | 73.47% | 31 |
| *Listeria* | 90.00% | 64.29% | 75.00% | 28 |
| *Arthrobacter* | 100.00% | 50.00% | 66.67% | 26 |
| *Caulobacter* | 100.00% | 64.00% | 78.05% | 25 |
| *Clostridium* | 66.67% | 16.67% | 26.67% | 24 |
| *Lactobacillus* | 50.00% | 4.35% | 8.00% | 23 |
| *Prevotella* | 100.00% | 83.33% | 90.91% | 18 |
| *Achromobacter* | 100.00% | 11.76% | 21.05% | 17 |
| *Pseudoalteromonas* | 100.00% | 11.76% | 21.05% | 17 |
| *Pantoea* | 0.00% | 0.00% | 0.00% | 16 |
| *Providencia* | 66.67% | 12.50% | 21.05% | 16 |
| *Helicobacter* | 100.00% | 87.50% | 93.33% | 16 |
| *Rhodococcus* | 0.00% | 0.00% | 0.00% | 13 |
| *Halorubrum* | 30.77% | 30.77% | 30.77% | 13 |
| *Agrobacterium* | 100.00% | 33.33% | 50.00% | 12 |
| *Shewanella* | 0.00% | 0.00% | 0.00% | 11 |
| *Leuconostoc* | 100.00% | 40.00% | 57.14% | 10 |
| *Kosakonia* | 0.00% | 0.00% | 0.00% | 10 |
| *Edwardsiella* | 0.00% | 0.00% | 0.00% | 10 |
| *Brevundimonas* | 100.00% | 20.00% | 33.33% | 10 |
| Others | 23.25% | 97.06% | 37.51% | 986 |

**S6 Table.**
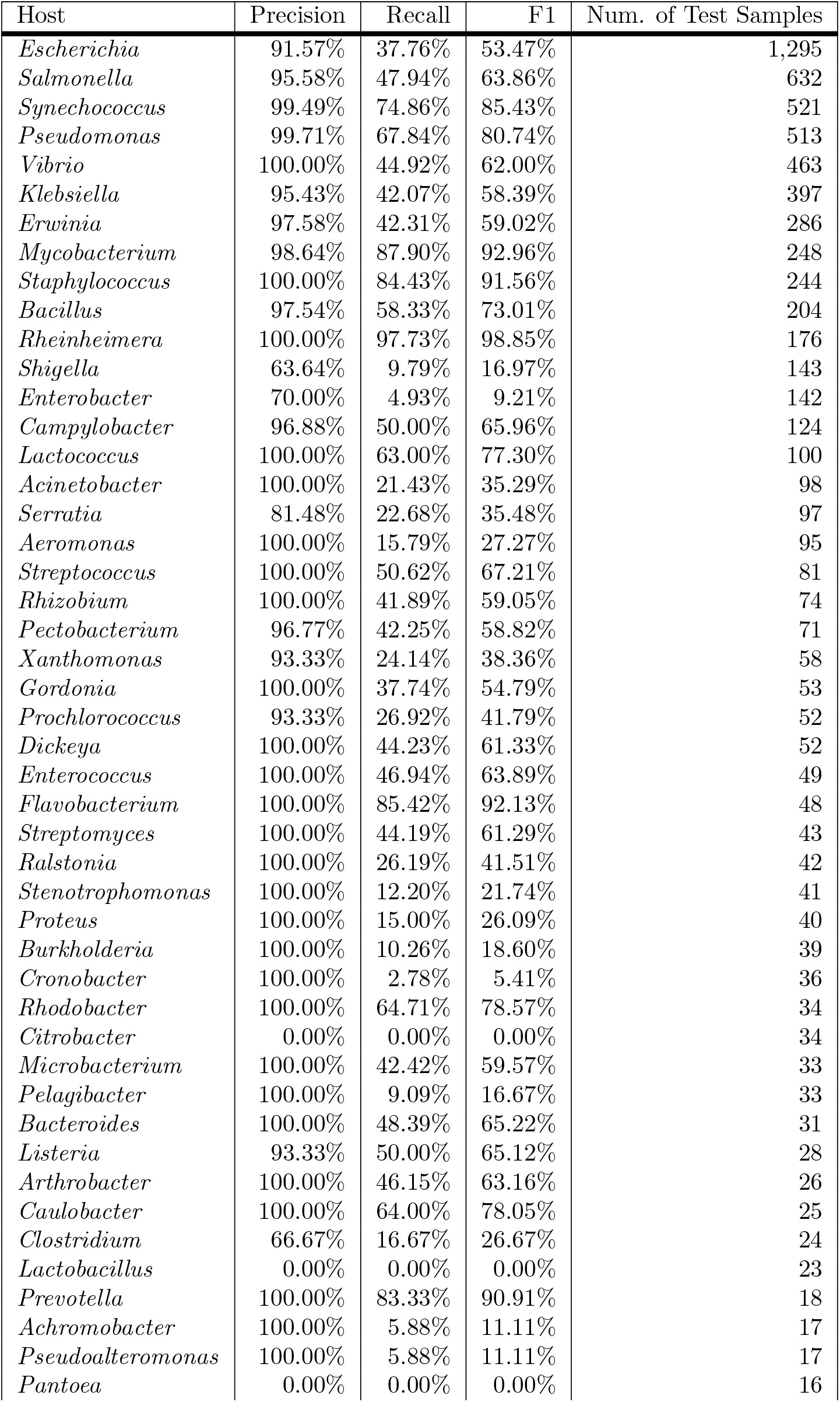
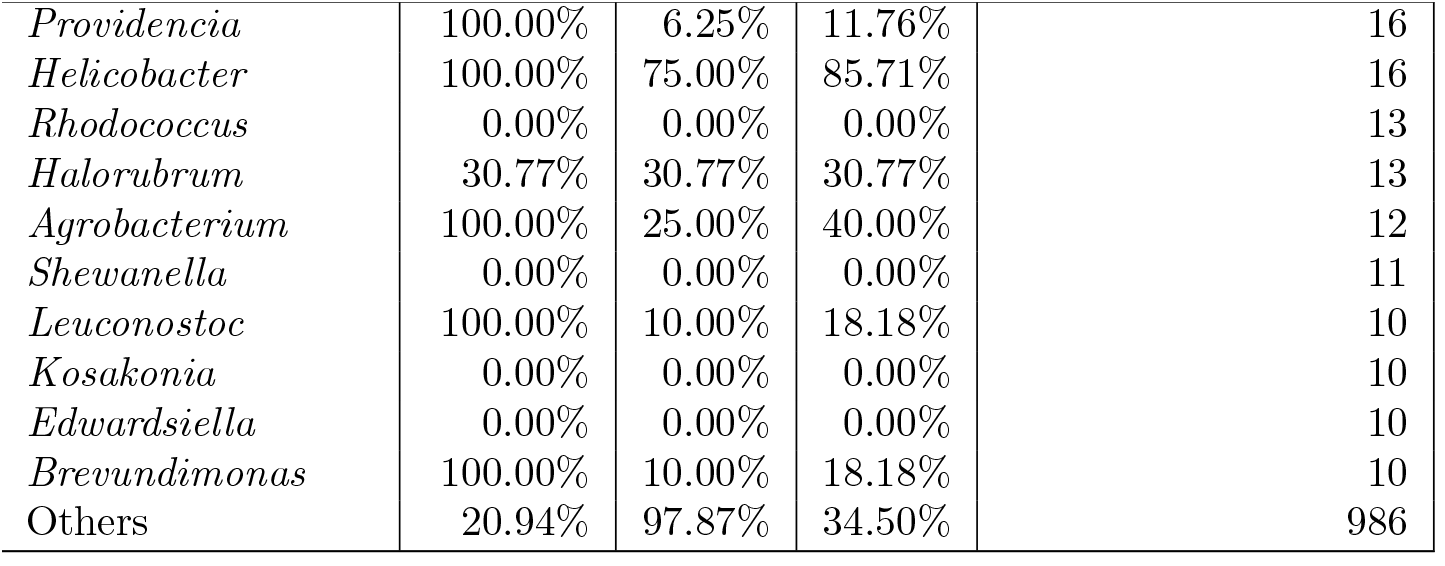
Per-class evaluation results of using ProtT5 embeddings at confidence threshold *k* = 70%.

| Host | Precision | Recall | F1 | Num. of Test Samples |
| --- | --- | --- | --- | --- |
| *Escherichia* | 91.57% | 37.76% | 53.47% | 1,295 |
| *Salmonella* | 95.58% | 47.94% | 63.86% | 632 |
| *Synechococcus* | 99.49% | 74.86% | 85.43% | 521 |
| *Pseudomonas* | 99.71% | 67.84% | 80.74% | 513 |
| *Vibrio* | 100.00% | 44.92% | 62.00% | 463 |
| *Klebsiella* | 95.43% | 42.07% | 58.39% | 397 |
| *Erwinia* | 97.58% | 42.31% | 59.02% | 286 |
| *Mycobacterium* | 98.64% | 87.90% | 92.96% | 248 |
| *Staphylococcus* | 100.00% | 84.43% | 91.56% | 244 |
| *Bacillus* | 97.54% | 58.33% | 73.01% | 204 |
| *Rheinheimera* | 100.00% | 97.73% | 98.85% | 176 |
| *Shigella* | 63.64% | 9.79% | 16.97% | 143 |
| *Enterobacter* | 70.00% | 4.93% | 9.21% | 142 |
| *Campylobacter* | 96.88% | 50.00% | 65.96% | 124 |
| *Lactococcus* | 100.00% | 63.00% | 77.30% | 100 |
| *Acinetobacter* | 100.00% | 21.43% | 35.29% | 98 |
| *Serratia* | 81.48% | 22.68% | 35.48% | 97 |
| *Aeromonas* | 100.00% | 15.79% | 27.27% | 95 |
| *Streptococcus* | 100.00% | 50.62% | 67.21% | 81 |
| *Rhizobium* | 100.00% | 41.89% | 59.05% | 74 |
| *Pectobacterium* | 96.77% | 42.25% | 58.82% | 71 |
| *Xanthomonas* | 93.33% | 24.14% | 38.36% | 58 |
| *Gordonia* | 100.00% | 37.74% | 54.79% | 53 |
| *Prochlorococcus* | 93.33% | 26.92% | 41.79% | 52 |
| *Dickeya* | 100.00% | 44.23% | 61.33% | 52 |
| *Enterococcus* | 100.00% | 46.94% | 63.89% | 49 |
| *Flavobacterium* | 100.00% | 85.42% | 92.13% | 48 |
| *Streptomyces* | 100.00% | 44.19% | 61.29% | 43 |
| *Ralstonia* | 100.00% | 26.19% | 41.51% | 42 |
| *Stenotrophomonas* | 100.00% | 12.20% | 21.74% | 41 |
| *Proteus* | 100.00% | 15.00% | 26.09% | 40 |
| *Burkholderia* | 100.00% | 10.26% | 18.60% | 39 |
| *Cronobacter* | 100.00% | 2.78% | 5.41% | 36 |
| *Rhodobacter* | 100.00% | 64.71% | 78.57% | 34 |
| *Citrobacter* | 0.00% | 0.00% | 0.00% | 34 |
| *Microbacterium* | 100.00% | 42.42% | 59.57% | 33 |
| *Pelagibacter* | 100.00% | 9.09% | 16.67% | 33 |
| *Bacteroides* | 100.00% | 48.39% | 65.22% | 31 |
| *Listeria* | 93.33% | 50.00% | 65.12% | 28 |
| *Arthrobacter* | 100.00% | 46.15% | 63.16% | 26 |
| *Caulobacter* | 100.00% | 64.00% | 78.05% | 25 |
| *Clostridium* | 66.67% | 16.67% | 26.67% | 24 |
| *Lactobacillus* | 0.00% | 0.00% | 0.00% | 23 |
| *Prevotella* | 100.00% | 83.33% | 90.91% | 18 |
| *Achromobacter* | 100.00% | 5.88% | 11.11% | 17 |
| *Pseudoalteromonas* | 100.00% | 5.88% | 11.11% | 17 |
| *Pantoea* | 0.00% | 0.00% | 0.00% | 16 |
| Continued on the next page |  |  |  |  |
| *Providencia* | 100.00% | 6.25% | 11.76% | 16 |
| *Helicobacter* | 100.00% | 75.00% | 85.71% | 16 |
| *Rhodococcus* | 0.00% | 0.00% | 0.00% | 13 |
| *Halorubrum* | 30.77% | 30.77% | 30.77% | 13 |
| *Agrobacterium* | 100.00% | 25.00% | 40.00% | 12 |
| *Shewanella* | 0.00% | 0.00% | 0.00% | 11 |
| *Leuconostoc* | 100.00% | 10.00% | 18.18% | 10 |
| *Kosakonia* | 0.00% | 0.00% | 0.00% | 10 |
| *Edwardsiella* | 0.00% | 0.00% | 0.00% | 10 |
| *Brevundimonas* | 100.00% | 10.00% | 18.18% | 10 |
| Others | 20.94% | 97.87% | 34.50% | 986 |

**S7 Table.**
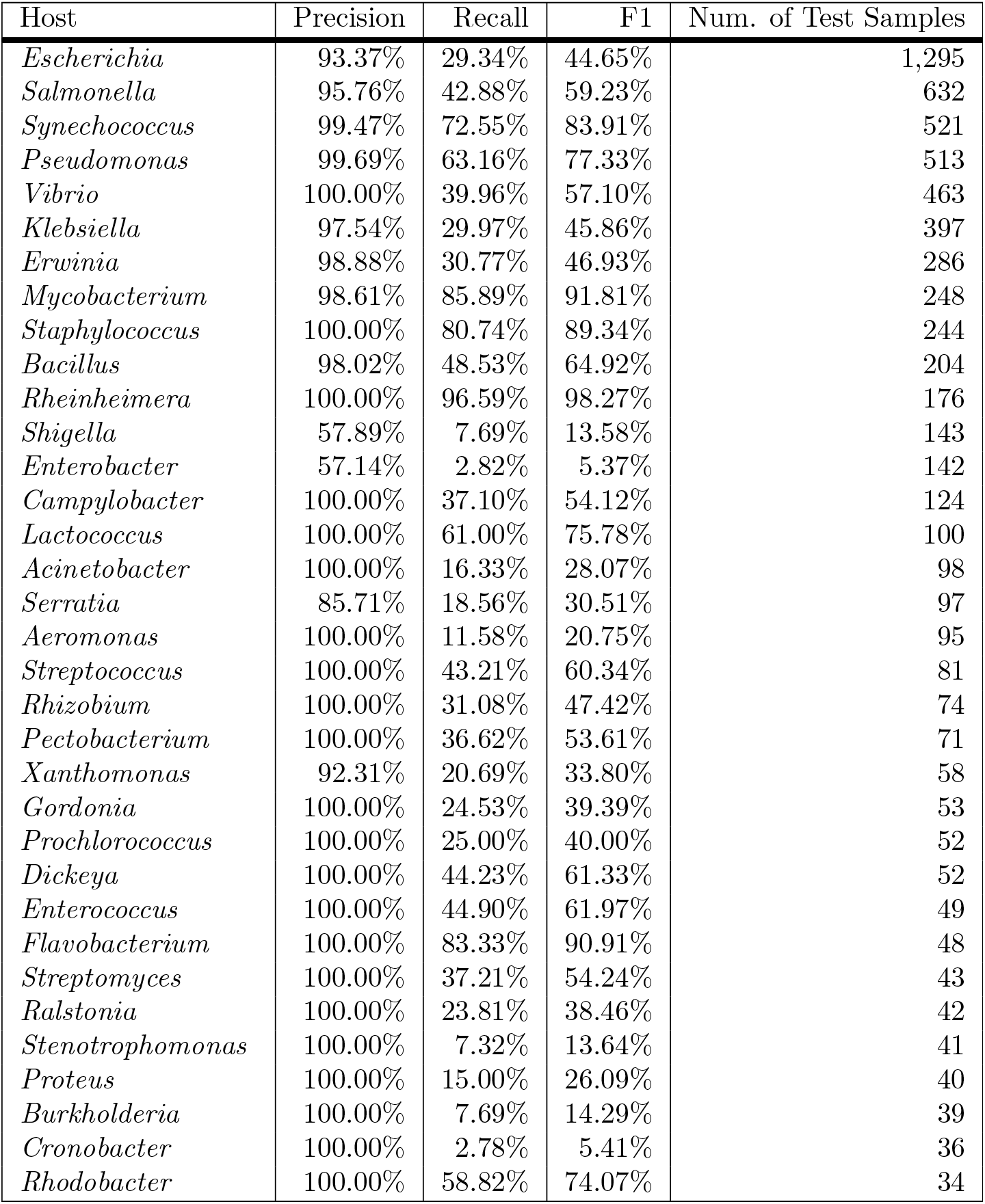
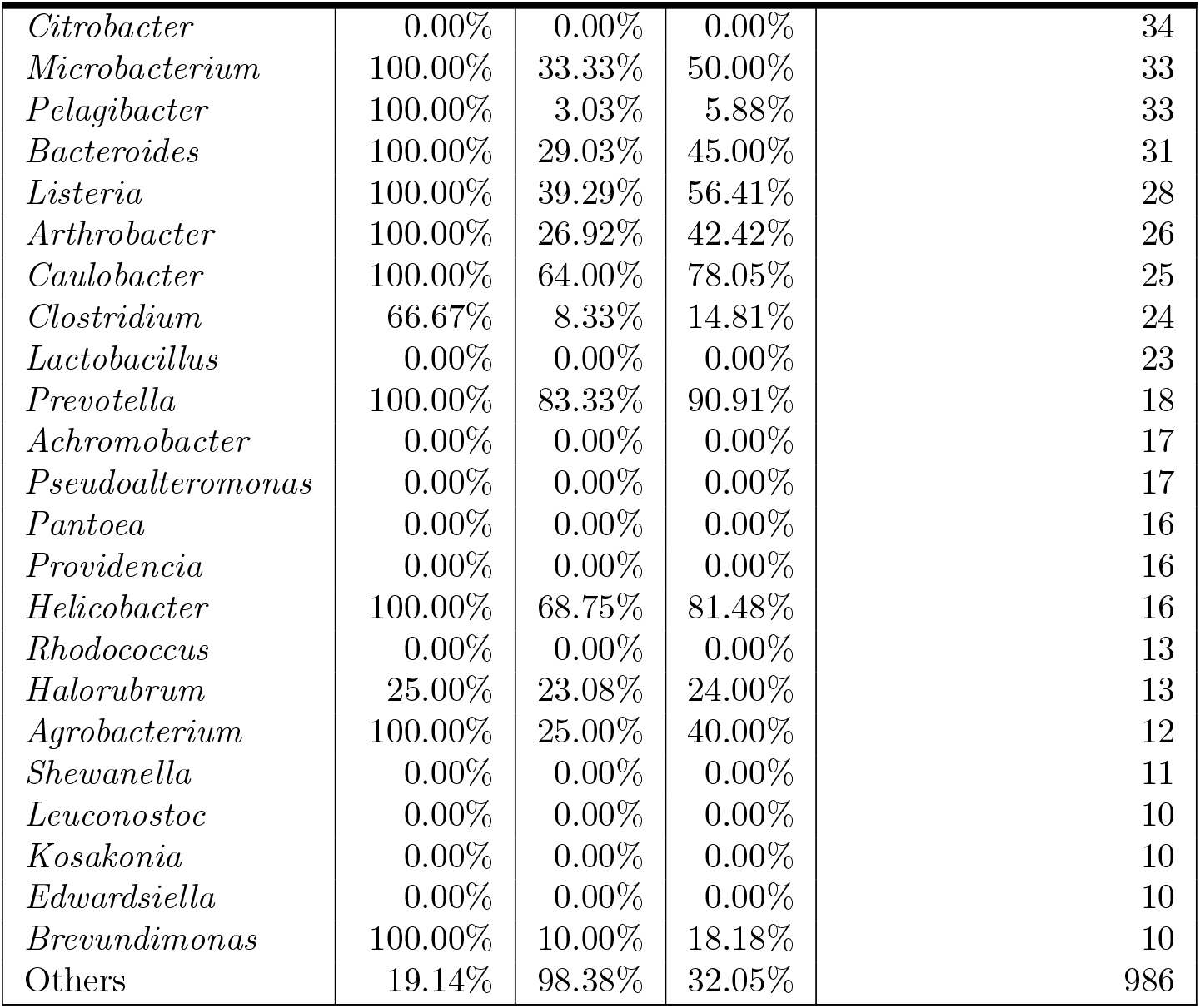
Per-class evaluation results of using ProtT5 embeddings at confidence threshold *k* = 80%.

| Host | Precision | Recall | F1 | Num. of Test Samples |
| --- | --- | --- | --- | --- |
| *Escherichia* | 93.37% | 29.34% | 44.65% | 1,295 |
| *Salmonella* | 95.76% | 42.88% | 59.23% | 632 |
| *Synechococcus* | 99.47% | 72.55% | 83.91% | 521 |
| *Pseudomonas* | 99.69% | 63.16% | 77.33% | 513 |
| *Vibrio* | 100.00% | 39.96% | 57.10% | 463 |
| *Klebsiella* | 97.54% | 29.97% | 45.86% | 397 |
| *Erwinia* | 98.88% | 30.77% | 46.93% | 286 |
| *Mycobacterium* | 98.61% | 85.89% | 91.81% | 248 |
| *Staphylococcus* | 100.00% | 80.74% | 89.34% | 244 |
| *Bacillus* | 98.02% | 48.53% | 64.92% | 204 |
| *Rheinheimera* | 100.00% | 96.59% | 98.27% | 176 |
| *Shigella* | 57.89% | 7.69% | 13.58% | 143 |
| *Enterobacter* | 57.14% | 2.82% | 5.37% | 142 |
| *Campylobacter* | 100.00% | 37.10% | 54.12% | 124 |
| *Lactococcus* | 100.00% | 61.00% | 75.78% | 100 |
| *Acinetobacter* | 100.00% | 16.33% | 28.07% | 98 |
| *Serratia* | 85.71% | 18.56% | 30.51% | 97 |
| *Aeromonas* | 100.00% | 11.58% | 20.75% | 95 |
| *Streptococcus* | 100.00% | 43.21% | 60.34% | 81 |
| *Rhizobium* | 100.00% | 31.08% | 47.42% | 74 |
| *Pectobacterium* | 100.00% | 36.62% | 53.61% | 71 |
| *Xanthomonas* | 92.31% | 20.69% | 33.80% | 58 |
| *Gordonia* | 100.00% | 24.53% | 39.39% | 53 |
| *Prochlorococcus* | 100.00% | 25.00% | 40.00% | 52 |
| *Dickeya* | 100.00% | 44.23% | 61.33% | 52 |
| *Enterococcus* | 100.00% | 44.90% | 61.97% | 49 |
| *Flavobacterium* | 100.00% | 83.33% | 90.91% | 48 |
| *Streptomyces* | 100.00% | 37.21% | 54.24% | 43 |
| *Ralstonia* | 100.00% | 23.81% | 38.46% | 42 |
| *Stenotrophomonas* | 100.00% | 7.32% | 13.64% | 41 |
| *Proteus* | 100.00% | 15.00% | 26.09% | 40 |
| *Burkholderia* | 100.00% | 7.69% | 14.29% | 39 |
| *Cronobacter* | 100.00% | 2.78% | 5.41% | 36 |
| *Rhodobacter* | 100.00% | 58.82% | 74.07% | 34 |
| *Citrobacter* | 0.00% | 0.00% | 0.00% | 34 |
| *Microbacterium* | 100.00% | 33.33% | 50.00% | 33 |
| *Pelagibacter* | 100.00% | 3.03% | 5.88% | 33 |
| *Bacteroides* | 100.00% | 29.03% | 45.00% | 31 |
| *Listeria* | 100.00% | 39.29% | 56.41% | 28 |
| *Arthrobacter* | 100.00% | 26.92% | 42.42% | 26 |
| *Caulobacter* | 100.00% | 64.00% | 78.05% | 25 |
| *Clostridium* | 66.67% | 8.33% | 14.81% | 24 |
| *Lactobacillus* | 0.00% | 0.00% | 0.00% | 23 |
| *Prevotella* | 100.00% | 83.33% | 90.91% | 18 |
| *Achromobacter* | 0.00% | 0.00% | 0.00% | 17 |
| *Pseudoalteromonas* | 0.00% | 0.00% | 0.00% | 17 |
| *Pantoea* | 0.00% | 0.00% | 0.00% | 16 |
| *Providencia* | 0.00% | 0.00% | 0.00% | 16 |
| *Helicobacter* | 100.00% | 68.75% | 81.48% | 16 |
| *Rhodococcus* | 0.00% | 0.00% | 0.00% | 13 |
| *Halorubrum* | 25.00% | 23.08% | 24.00% | 13 |
| *Agrobacterium* | 100.00% | 25.00% | 40.00% | 12 |
| *Shewanella* | 0.00% | 0.00% | 0.00% | 11 |
| *Leuconostoc* | 0.00% | 0.00% | 0.00% | 10 |
| *Kosakonia* | 0.00% | 0.00% | 0.00% | 10 |
| *Edwardsiella* | 0.00% | 0.00% | 0.00% | 10 |
| *Brevundimonas* | 100.00% | 10.00% | 18.18% | 10 |
| Others | 19.14% | 98.38% | 32.05% | 986 |

**S8 Table.**
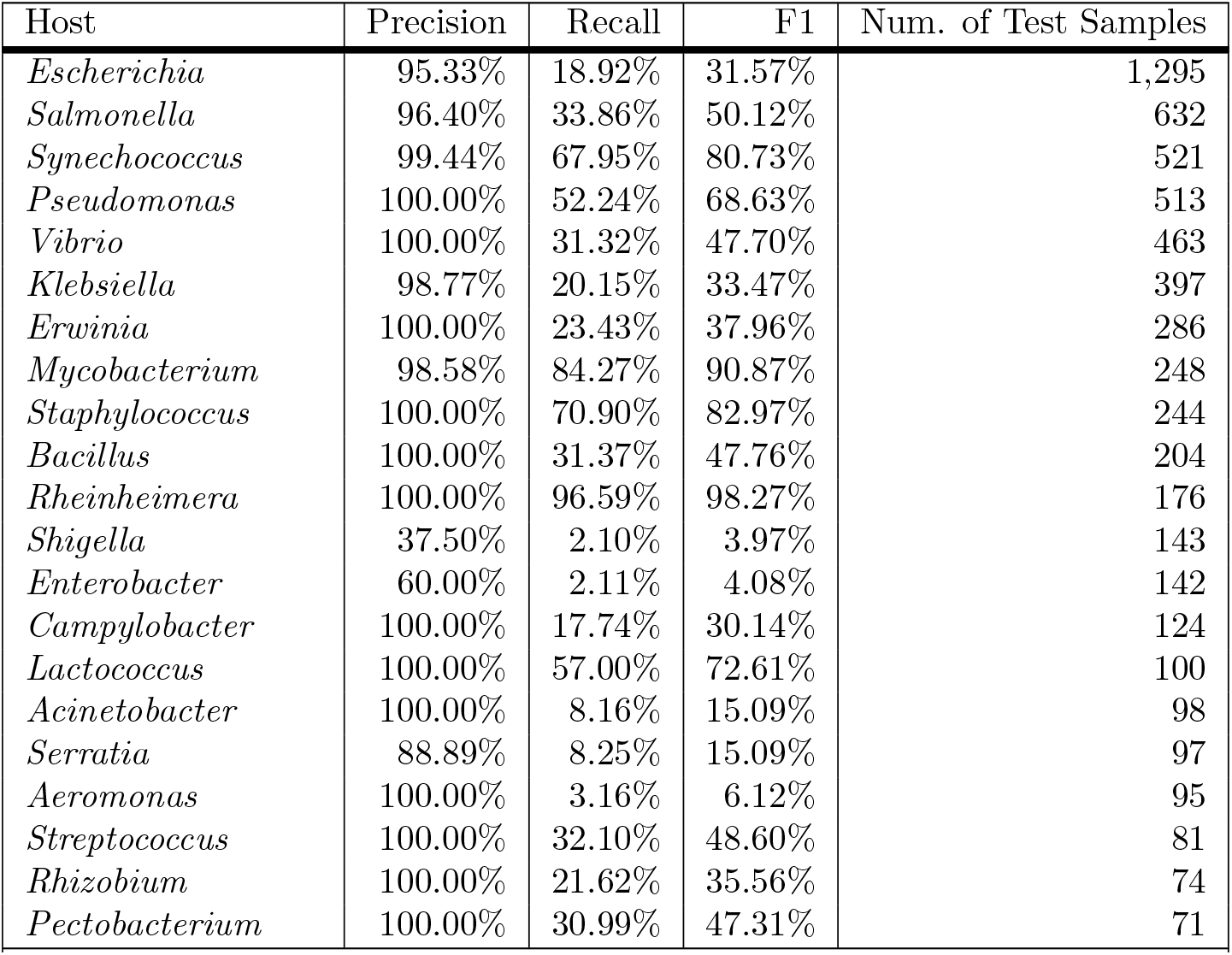
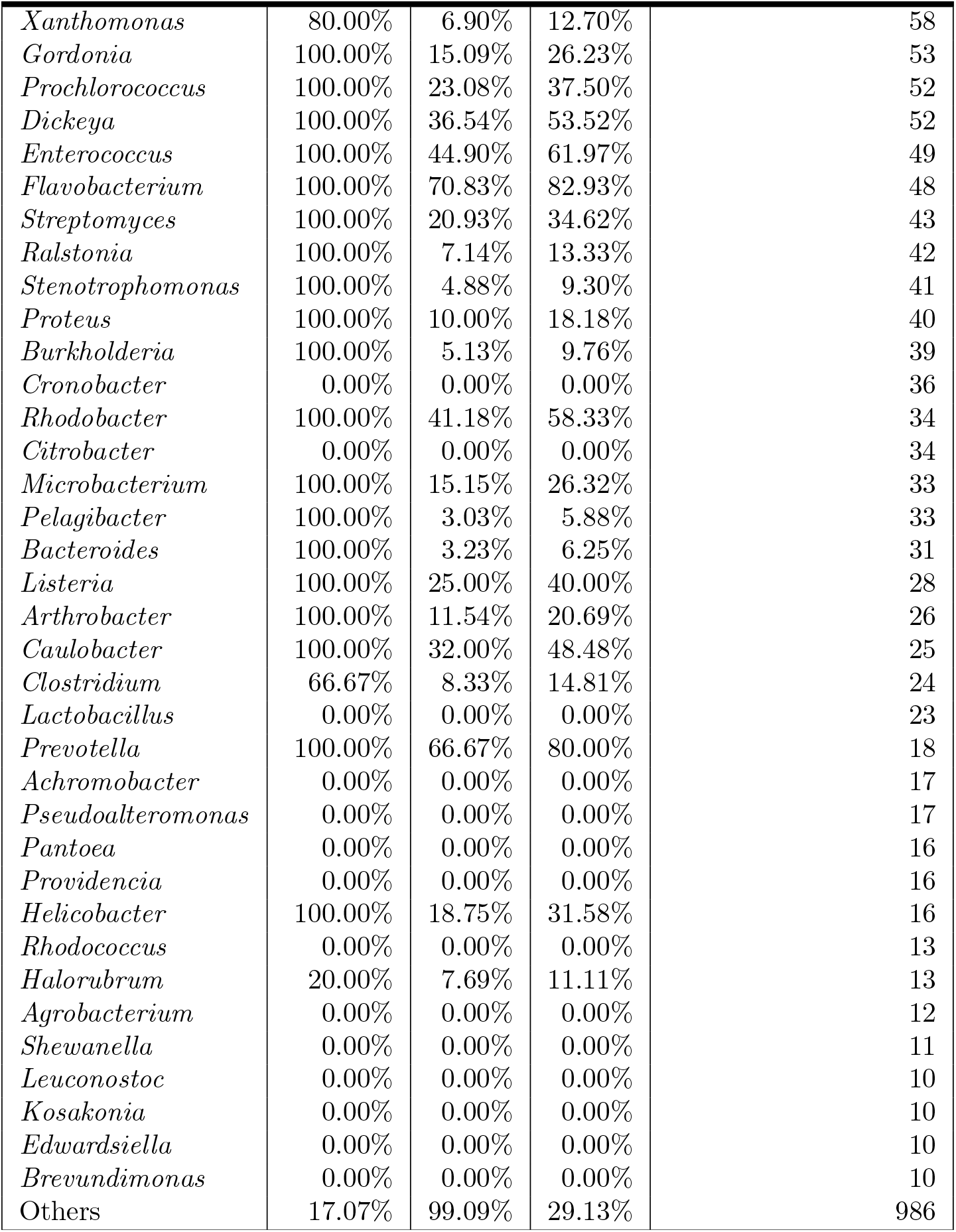
Per-class evaluation results of using ProtT5 embeddings at confidence threshold *k* = 90%.

| Host | Precision | Recall | F1 | Num. of Test Samples |
| --- | --- | --- | --- | --- |
| *Escherichia* | 95.33% | 18.92% | 31.57% | 1,295 |
| *Salmonella* | 96.40% | 33.86% | 50.12% | 632 |
| *Synechococcus* | 99.44% | 67.95% | 80.73% | 521 |
| *Pseudomonas* | 100.00% | 52.24% | 68.63% | 513 |
| *Vibrio* | 100.00% | 31.32% | 47.70% | 463 |
| *Klebsiella* | 98.77% | 20.15% | 33.47% | 397 |
| *Erwinia* | 100.00% | 23.43% | 37.96% | 286 |
| *Mycobacterium* | 98.58% | 84.27% | 90.87% | 248 |
| *Staphylococcus* | 100.00% | 70.90% | 82.97% | 244 |
| *Bacillus* | 100.00% | 31.37% | 47.76% | 204 |
| *Rheinheimera* | 100.00% | 96.59% | 98.27% | 176 |
| *Shigella* | 37.50% | 2.10% | 3.97% | 143 |
| *Enterobacter* | 60.00% | 2.11% | 4.08% | 142 |
| *Campylobacter* | 100.00% | 17.74% | 30.14% | 124 |
| *Lactococcus* | 100.00% | 57.00% | 72.61% | 100 |
| *Acinetobacter* | 100.00% | 8.16% | 15.09% | 98 |
| *Serratia* | 88.89% | 8.25% | 15.09% | 97 |
| *Aeromonas* | 100.00% | 3.16% | 6.12% | 95 |
| *Streptococcus* | 100.00% | 32.10% | 48.60% | 81 |
| *Rhizobium* | 100.00% | 21.62% | 35.56% | 74 |
| *Pectobacterium* | 100.00% | 30.99% | 47.31% | 71 |
| *Xanthomonas* | 80.00% | 6.90% | 12.70% | 58 |
| *Gordonia* | 100.00% | 15.09% | 26.23% | 53 |
| *Prochlorococcus* | 100.00% | 23.08% | 37.50% | 52 |
| *Dickeya* | 100.00% | 36.54% | 53.52% | 52 |
| *Enterococcus* | 100.00% | 44.90% | 61.97% | 49 |
| *Flavobacterium* | 100.00% | 70.83% | 82.93% | 48 |
| *Streptomyces* | 100.00% | 20.93% | 34.62% | 43 |
| *Ralstonia* | 100.00% | 7.14% | 13.33% | 42 |
| *Stenotrophomonas* | 100.00% | 4.88% | 9.30% | 41 |
| *Proteus* | 100.00% | 10.00% | 18.18% | 40 |
| *Burkholderia* | 100.00% | 5.13% | 9.76% | 39 |
| *Cronobacter* | 0.00% | 0.00% | 0.00% | 36 |
| *Rhodobacter* | 100.00% | 41.18% | 58.33% | 34 |
| *Citrobacter* | 0.00% | 0.00% | 0.00% | 34 |
| *Microbacterium* | 100.00% | 15.15% | 26.32% | 33 |
| *Pelagibacter* | 100.00% | 3.03% | 5.88% | 33 |
| *Bacteroides* | 100.00% | 3.23% | 6.25% | 31 |
| *Listeria* | 100.00% | 25.00% | 40.00% | 28 |
| *Arthrobacter* | 100.00% | 11.54% | 20.69% | 26 |
| *Caulobacter* | 100.00% | 32.00% | 48.48% | 25 |
| *Clostridium* | 66.67% | 8.33% | 14.81% | 24 |
| *Lactobacillus* | 0.00% | 0.00% | 0.00% | 23 |
| *Prevotella* | 100.00% | 66.67% | 80.00% | 18 |
| *Achromobacter* | 0.00% | 0.00% | 0.00% | 17 |
| *Pseudoalteromonas* | 0.00% | 0.00% | 0.00% | 17 |
| *Pantoea* | 0.00% | 0.00% | 0.00% | 16 |
| *Providencia* | 0.00% | 0.00% | 0.00% | 16 |
| *Helicobacter* | 100.00% | 18.75% | 31.58% | 16 |
| *Rhodococcus* | 0.00% | 0.00% | 0.00% | 13 |
| *Halorubrum* | 20.00% | 7.69% | 11.11% | 13 |
| *Agrobacterium* | 0.00% | 0.00% | 0.00% | 12 |
| *Shewanella* | 0.00% | 0.00% | 0.00% | 11 |
| *Leuconostoc* | 0.00% | 0.00% | 0.00% | 10 |
| *Kosakonia* | 0.00% | 0.00% | 0.00% | 10 |
| *Edwardsiella* | 0.00% | 0.00% | 0.00% | 10 |
| *Brevundimonas* | 0.00% | 0.00% | 0.00% | 10 |
| Others | 17.07% | 99.09% | 29.13% | 986 |

**S9 Table.**
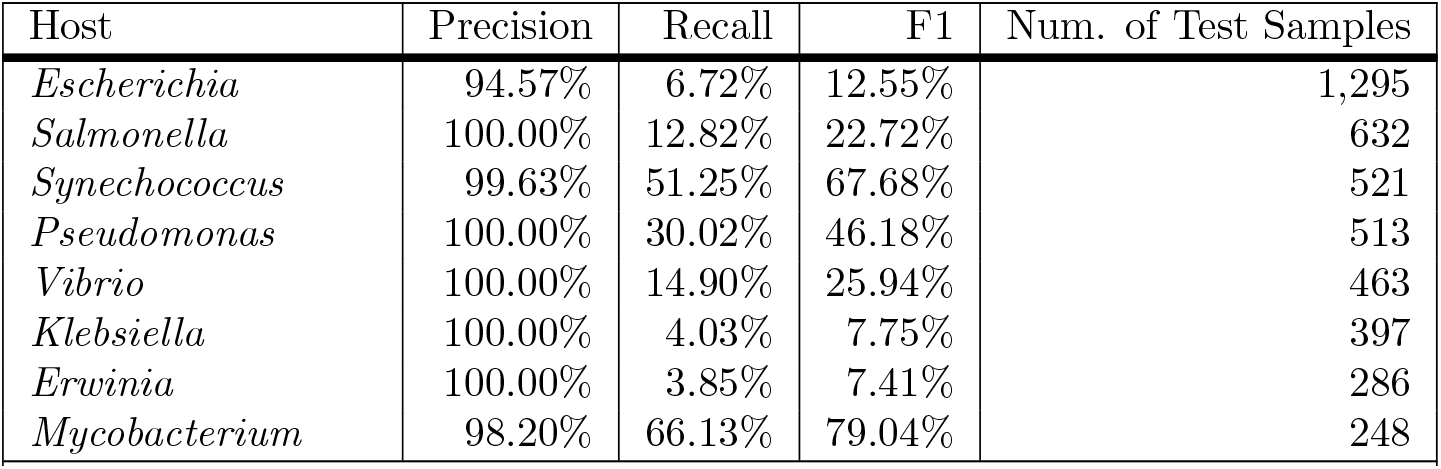
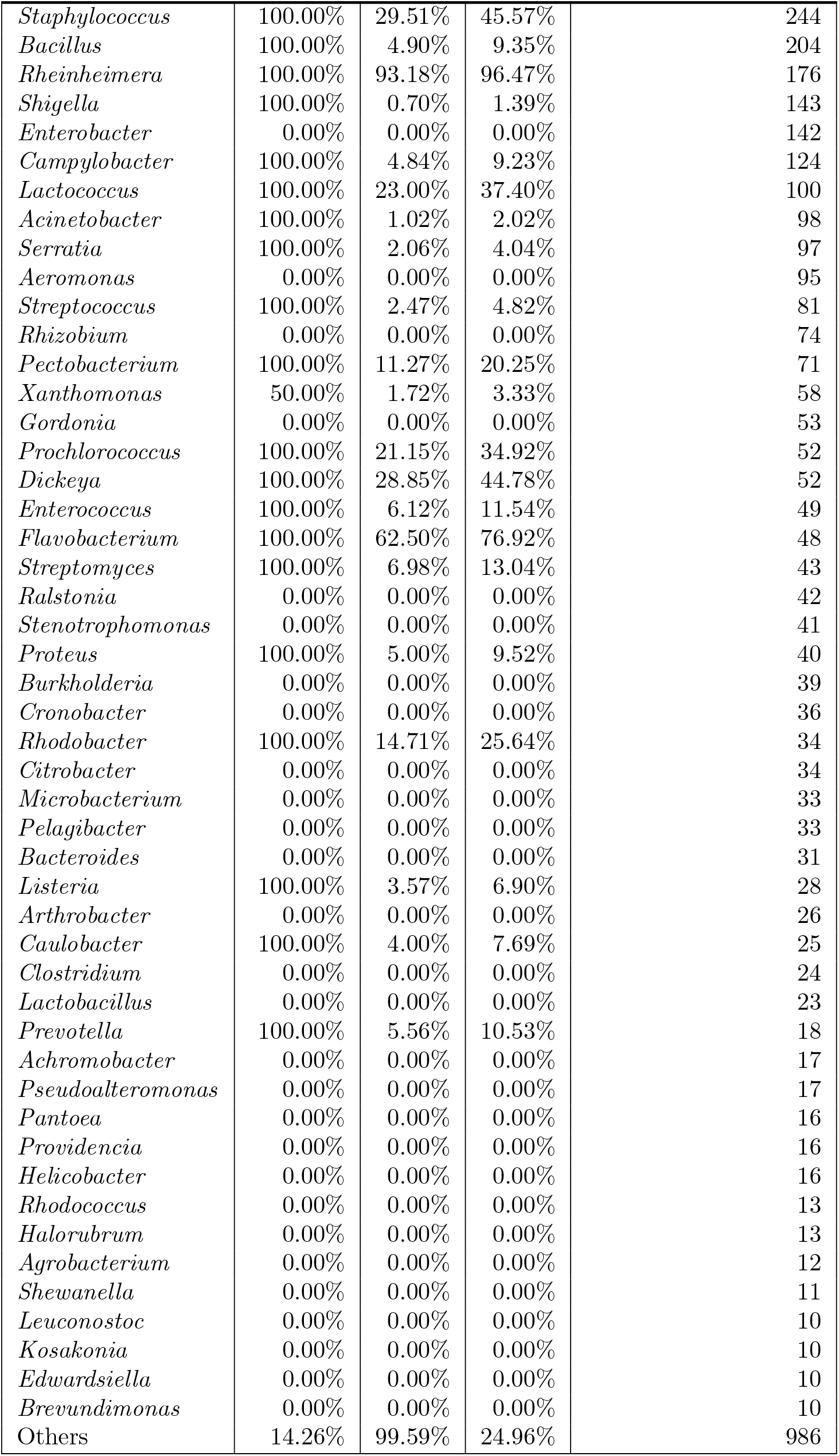
Per-class evaluation results of using ProtT5 embeddings at confidence threshold *k* = 100%.

| Host | Precision | Recall | F1 | Num. of Test Samples |
| --- | --- | --- | --- | --- |
| *Escherichia* | 94.57% | 6.72% | 12.55% | 1,295 |
| *Salmonella* | 100.00% | 12.82% | 22.72% | 632 |
| *Synechococcus* | 99.63% | 51.25% | 67.68% | 521 |
| *Pseudomonas* | 100.00% | 30.02% | 46.18% | 513 |
| *Vibrio* | 100.00% | 14.90% | 25.94% | 463 |
| *Klebsiella* | 100.00% | 4.03% | 7.75% | 397 |
| *Erwinia* | 100.00% | 3.85% | 7.41% | 286 |
| *Mycobacterium* | 98.20% | 66.13% | 79.04% | 248 |
| *Staphylococcus* | 100.00% | 29.51% | 45.57% | 244 |
| *Bacillus* | 100.00% | 4.90% | 9.35% | 204 |
| *Rheinheimera* | 100.00% | 93.18% | 96.47% | 176 |
| *Shigella* | 100.00% | 0.70% | 1.39% | 143 |
| *Enterobacter* | 0.00% | 0.00% | 0.00% | 142 |
| *Campylobacter* | 100.00% | 4.84% | 9.23% | 124 |
| *Lactococcus* | 100.00% | 23.00% | 37.40% | 100 |
| *Acinetobacter* | 100.00% | 1.02% | 2.02% | 98 |
| *Serratia* | 100.00% | 2.06% | 4.04% | 97 |
| *Aeromonas* | 0.00% | 0.00% | 0.00% | 95 |
| *Streptococcus* | 100.00% | 2.47% | 4.82% | 81 |
| *Rhizobium* | 0.00% | 0.00% | 0.00% | 74 |
| *Pectobacterium* | 100.00% | 11.27% | 20.25% | 71 |
| *Xanthomonas* | 50.00% | 1.72% | 3.33% | 58 |
| *Gordonia* | 0.00% | 0.00% | 0.00% | 53 |
| *Prochlorococcus* | 100.00% | 21.15% | 34.92% | 52 |
| *Dickeya* | 100.00% | 28.85% | 44.78% | 52 |
| *Enterococcus* | 100.00% | 6.12% | 11.54% | 49 |
| *Flavobacterium* | 100.00% | 62.50% | 76.92% | 48 |
| *Streptomyces* | 100.00% | 6.98% | 13.04% | 43 |
| *Ralstonia* | 0.00% | 0.00% | 0.00% | 42 |
| *Stenotrophomonas* | 0.00% | 0.00% | 0.00% | 41 |
| *Proteus* | 100.00% | 5.00% | 9.52% | 40 |
| *Burkholderia* | 0.00% | 0.00% | 0.00% | 39 |
| *Cronobacter* | 0.00% | 0.00% | 0.00% | 36 |
| *Rhodobacter* | 100.00% | 14.71% | 25.64% | 34 |
| *Citrobacter* | 0.00% | 0.00% | 0.00% | 34 |
| *Microbacterium* | 0.00% | 0.00% | 0.00% | 33 |
| *Pelagibacter* | 0.00% | 0.00% | 0.00% | 33 |
| *Bacteroides* | 0.00% | 0.00% | 0.00% | 31 |
| *Listeria* | 100.00% | 3.57% | 6.90% | 28 |
| *Arthrobacter* | 0.00% | 0.00% | 0.00% | 26 |
| *Caulobacter* | 100.00% | 4.00% | 7.69% | 25 |
| *Clostridium* | 0.00% | 0.00% | 0.00% | 24 |
| *Lactobacillus* | 0.00% | 0.00% | 0.00% | 23 |
| *Prevotella* | 100.00% | 5.56% | 10.53% | 18 |
| *Achromobacter* | 0.00% | 0.00% | 0.00% | 17 |
| *Pseudoalteromonas* | 0.00% | 0.00% | 0.00% | 17 |
| *Pantoea* | 0.00% | 0.00% | 0.00% | 16 |
| *Providencia* | 0.00% | 0.00% | 0.00% | 16 |
| *Helicobacter* | 0.00% | 0.00% | 0.00% | 16 |
| *Rhodococcus* | 0.00% | 0.00% | 0.00% | 13 |
| *Halorubrum* | 0.00% | 0.00% | 0.00% | 13 |
| *Agrobacterium* | 0.00% | 0.00% | 0.00% | 12 |
| *Shewanella* | 0.00% | 0.00% | 0.00% | 11 |
| *Leuconostoc* | 0.00% | 0.00% | 0.00% | 10 |
| *Kosakonia* | 0.00% | 0.00% | 0.00% | 10 |
| *Edwardsiella* | 0.00% | 0.00% | 0.00% | 10 |
| *Brevundimonas* | 0.00% | 0.00% | 0.00% | 10 |
| Others | 14.26% | 99.59% | 24.96% | 986 |

**S10 Table.** Weighted precision scores after integrating handcrafted sequence properties to the vector representations of the receptor-binding proteins (RBPs).

|  | 60% | 70% | 80% | 90% | 100% |
| --- | --- | --- | --- | --- | --- |
| ProtT5 | 85.43% | 84.98% | 84.32% | 83.51% | 77.23% |
| ProtT5 + A Frequency | 85.86% | 85.05% | 84.48% | 83.47% | 76.52% |
| ProtT5 + GC Content | 85.89% | 84.93% | 84.22% | 84.10% | 75.47% |
| ProtT5 + C Frequency | 85.44% | 84.80% | 84.43% | 83.79% | 77.00% |
| ProtT5 + TTA Frequency | 85.46% | 84.81% | 84.34% | 83.60% | 77.03% |
| ProtT5 + TTA Bias | 85.69% | 84.78% | 84.89% | 84.01% | 75.16% |

This table shows the weighted precision scores after combining the ProtT5 embeddings of the RBPs with the sequence properties that registered the highest Gini importance after training the phage-host interaction prediction tool by Boeckaerts *et al.* [15] on our dataset. *A Frequency* stands for A nucleotide frequency; *GC Content*, guanine-cytosine content; *C Frequency*, C nucleotide frequency; *TTA Frequency*, TTA codon frequency; and *TTA Bias*, TTA codon usage bias (TTA codes for the amino acid leucine). The header row refers to the confidence thresholds at which we evaluated model performance.

**S11 Table.**
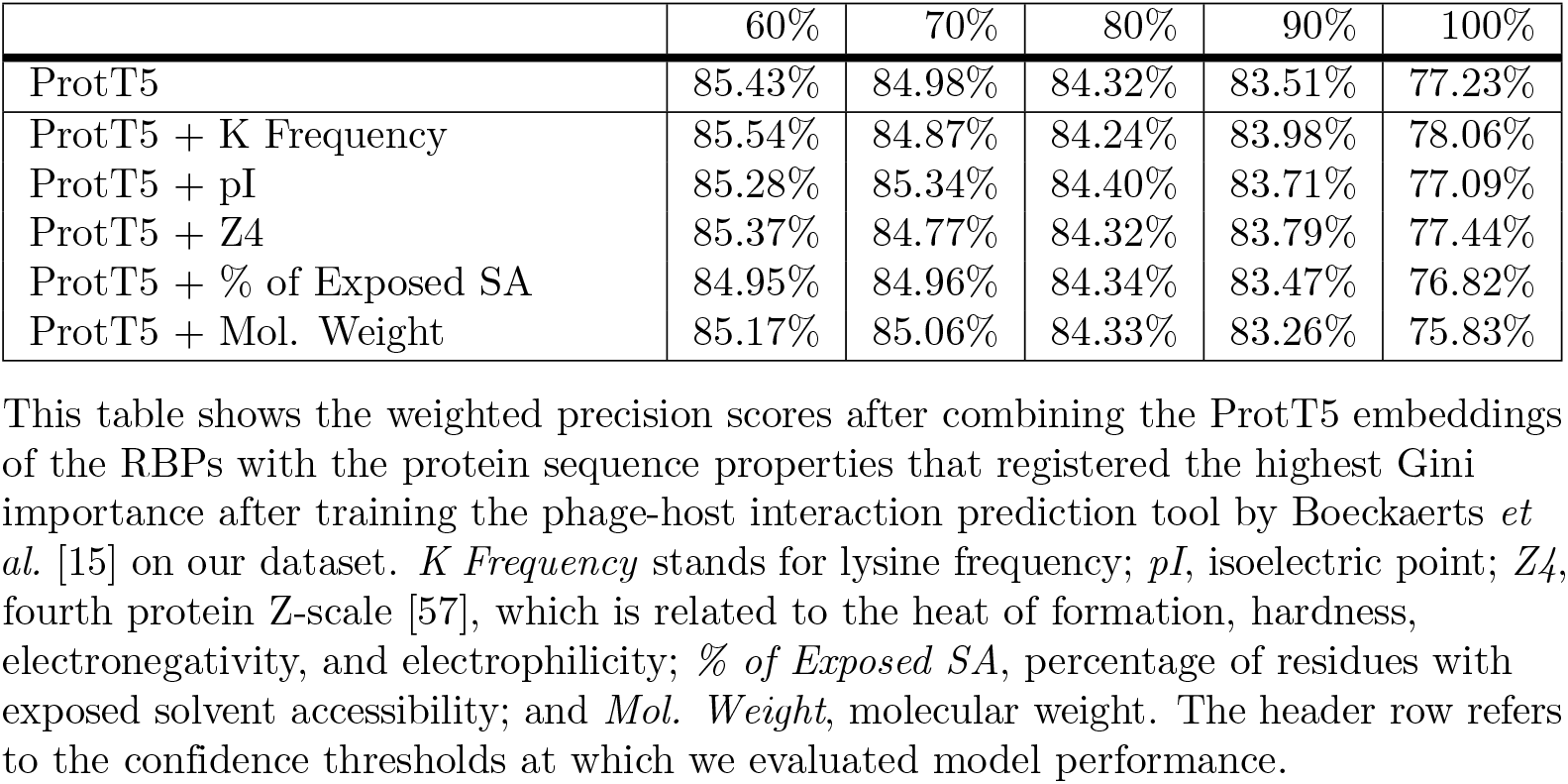
Weighted precision scores after integrating handcrafted protein sequence properties to the vector representations of the receptor-binding proteins (RBPs).

|  | 60% | 70% | 80% | 90% | 100% |
| --- | --- | --- | --- | --- | --- |
| ProtT5 | 85.43% | 84.98% | 84.32% | 83.51% | 77.23% |
| ProtT5 + K Frequency | 85.54% | 84.87% | 84.24% | 83.98% | 78.06% |
| ProtT5 + pI | 85.28% | 85.34% | 84.40% | 83.71% | 77.09% |
| ProtT5 + Z4 | 85.37% | 84.77% | 84.32% | 83.79% | 77.44% |
| ProtT5 + % of Exposed SA | 84.95% | 84.96% | 84.34% | 83.47% | 76.82% |
| ProtT5 + Mol. Weight | 85.17% | 85.06% | 84.33% | 83.26% | 75.83% |

This table shows the weighted precision scores after combining the ProtT5 embeddings of the RBPs with the protein sequence properties that registered the highest Gini importance after training the phage-host interaction prediction tool by Boeckaerts *et al.* [15] on our dataset. *K Frequency* stands for lysine frequency; *pI*, isoelectric point; *Z4*, fourth protein Z-scale [57], which is related to the heat of formation, hardness, electronegativity, and electrophilicity; *% of Exposed SA*, percentage of residues with exposed solvent accessibility; and *Mol. Weight*, molecular weight. The header row refers to the confidence thresholds at which we evaluated model performance.

**S12 Table.**
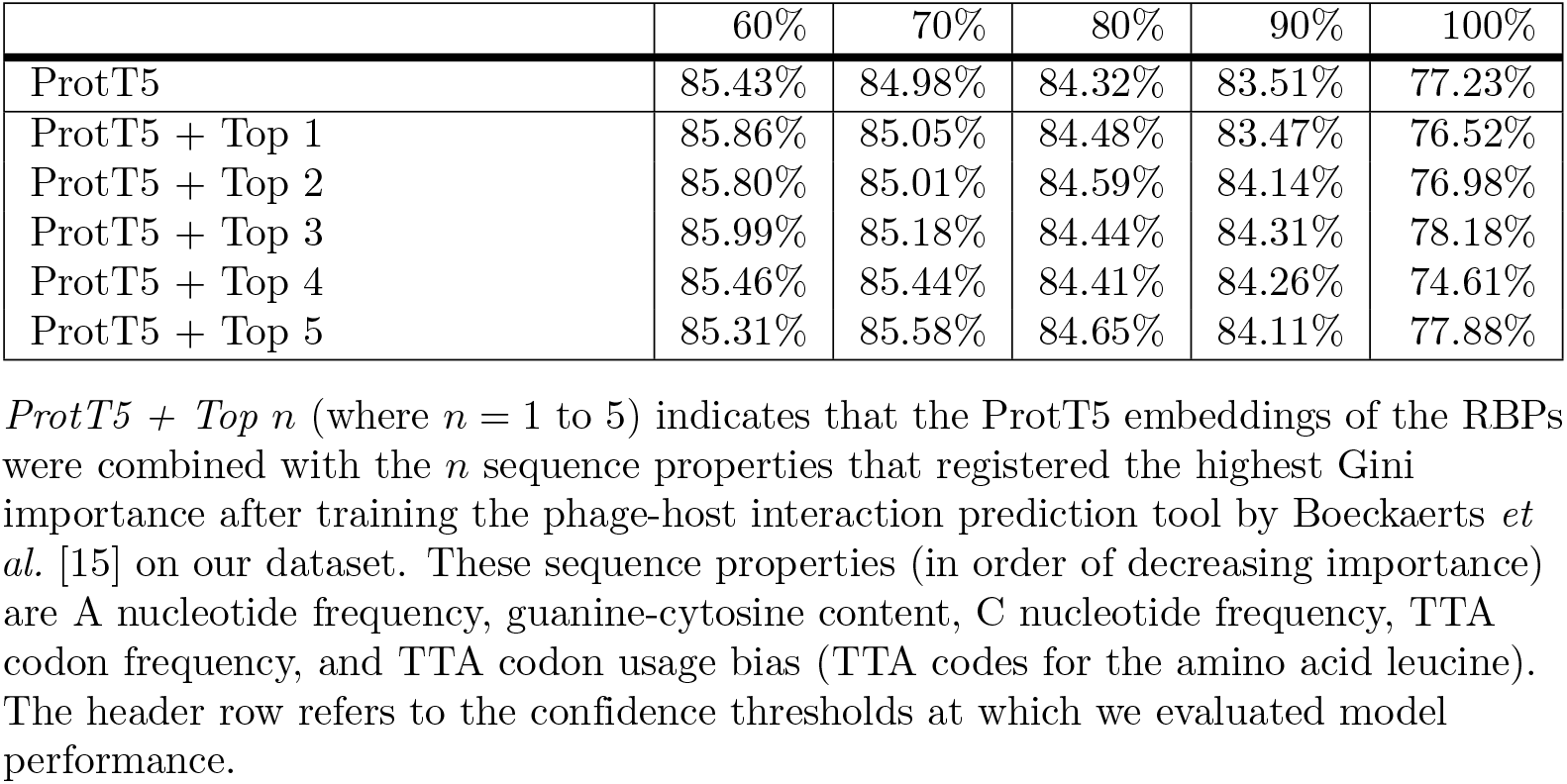
Weighted precision scores after integrating the top *n* handcrafted sequence properties to the vector representations of the receptor-binding proteins (RBPs).

**S13 Table.** Weighted precision scores after integrating the top *n* handcrafted protein sequence properties to the vector representations of the receptor-binding proteins (RBPs).

|  | 60% | 70% | 80% | 90% | 100% |
| --- | --- | --- | --- | --- | --- |
| ProtT5 | 85.43% | 84.98% | 84.32% | 83.51% | 77.23% |
| ProtT5 + Protein Top 1 | 85.54% | 84.87% | 84.24% | 83.98% | 78.06% |
| ProtT5 + Protein Top 2 | 85.53% | 84.88% | 84.29% | 83.24% | 75.05% |
| ProtT5 + Protein Top 3 | 85.19% | 85.32% | 84.12% | 83.90% | 78.62% |
| ProtT5 + Protein Top 4 | 85.18% | 84.98% | 83.99% | 83.86% | 77.62% |
| ProtT5 + Protein Top 5 | 85.42% | 85.29% | 84.51% | 83.83% | 75.81% |

*ProtT5 + Protein Top $n$* (where $n = 1$ to 5) indicates that the ProtT5 embeddings of the RBPs were combined with the $n$ protein sequence properties that registered the highest Gini importance after training the phage-host interaction prediction tool by Boeckaerts *et al.* [15] on our dataset. These sequence properties (in order of decreasing importance) are lysine frequency, isoelectric point, fourth protein Z-scale [57] (which is related to the heat of formation, hardness, electronegativity, and electrophilicity), percentage of residues with exposed solvent accessibility, and molecular weight. The header row refers to the confidence thresholds at which we evaluated model performance.

**S14 Table.** Weighted recall scores after integrating handcrafted sequence properties to the vector representations of the receptor-binding proteins (RBPs).

|  | 60% | 70% | 80% | 90% | 100% |
| --- | --- | --- | --- | --- | --- |
| ProtT5 | 59.15% | 53.72% | 48.57% | 41.03% | 27.16% |
| ProtT5 + A Frequency | 59.71% | 54.51% | 49.14% | 41.29% | 27.16% |
| ProtT5 + GC Content | 60.20% | 54.77% | 49.15% | 41.74% | 27.70% |
| ProtT5 + C Frequency | 59.91% | 54.62% | 48.98% | 41.38% | 27.24% |
| ProtT5 + TTA Frequency | 59.51% | 53.77% | 48.39% | 41.01% | 27.23% |
| ProtT5 + TTA Bias | 59.41% | 53.78% | 48.32% | 40.99% | 27.07% |

This table shows the weighted recall scores after combining the ProtT5 embeddings of the RBPs with the sequence properties that registered the highest Gini importance after training the phage-host interaction prediction tool by Boeckaerts *et al.* [15] on our dataset. *A Frequency* stands for A nucleotide frequency; *GC Content*, guanine-cytosine content; *C Frequency*, C nucleotide frequency; *TTA Frequency*, TTA codon frequency; and *TTA Bias*, TTA codon usage bias (TTA codes for the amino acid leucine). The header row refers to the confidence thresholds at which we evaluated model performance.

**S15 Table.** Weighted recall scores after integrating handcrafted protein sequence properties to the vector representations of the receptor-binding proteins (RBPs).

|  | 60% | 70% | 80% | 90% | 100% |
| --- | --- | --- | --- | --- | --- |
| ProtT5 | 59.15% | 53.72% | 48.57% | 41.03% | 27.16% |
| ProtT5 + K Frequency | 59.22% | 53.67% | 48.63% | 41.13% | 27.23% |
| ProtT5 + pI | 59.13% | 53.55% | 48.44% | 40.93% | 27.51% |
| ProtT5 + Z4 | 59.09% | 53.63% | 48.18% | 41.01% | 27.23% |
| ProtT5 + % of Exposed SA | 59.01% | 53.82% | 48.36% | 40.88% | 27.58% |
| ProtT5 + Mol. Weight | 59.22% | 53.87% | 48.46% | 40.86% | 27.34% |

This table shows the weighted recall scores after combining the ProtT5 embeddings of the RBPs with the protein sequence properties that registered the highest Gini importance after training the phage-host interaction prediction tool by Boeckaerts *et al.* [15] on our dataset. *K Frequency* stands for lysine frequency; *pI*, isoelectric point; *Z4*, fourth protein Z-scale [57], which is related to the heat of formation, hardness, electronegativity, and electrophilicity; *% of Exposed SA*, percentage of residues with exposed solvent accessibility; and *Mol. Weight*, molecular weight. The header row refers to the confidence thresholds at which we evaluated model performance.

**S16 Table.** Weighted recall scores after integrating the top *n* handcrafted sequence properties to the vector representations of the receptor-binding proteins (RBPs).

|  | 60% | 70% | 80% | 90% | 100% |
| --- | --- | --- | --- | --- | --- |
| ProtT5 | 59.15% | 53.72% | 48.57% | 41.03% | 27.16% |
| ProtT5 + Top 1 | 59.71% | 54.51% | 49.14% | 41.29% | 27.16% |
| ProtT5 + Top 2 | 60.55% | 54.95% | 49.22% | 41.51% | 27.37% |
| ProtT5 + Top 3 | 60.45% | 55.01% | 49.37% | 41.93% | 27.39% |
| ProtT5 + Top 4 | 60.31% | 55.04% | 49.25% | 41.67% | 26.81% |
| ProtT5 + Top 5 | 60.33% | 54.87% | 49.33% | 41.94% | 27.13% |

*ProtT5 + Top $n$* (where $n = 1$ to 5) indicates that the ProtT5 embeddings of the RBPs were combined with the $n$ sequence properties that registered the highest Gini importance after training the phage-host interaction prediction tool by Boeckaerts *et al.* [15] on our dataset. These sequence properties (in order of decreasing importance) are A nucleotide frequency, guanine-cytosine content, C nucleotide frequency, TTA codon frequency, and TTA codon usage bias (TTA codes for the amino acid leucine). The header row refers to the confidence thresholds at which we evaluated model performance.

**S17 Table.** Weighted recall scores after integrating the top *n* handcrafted protein sequence properties to the vector representations of the receptor-binding proteins (RBPs).

|  | 60% | 70% | 80% | 90% | 100% |
| --- | --- | --- | --- | --- | --- |
| ProtT5 | 59.15% | 53.72% | 48.57% | 41.03% | 27.16% |
| ProtT5 + Protein Top 1 | 59.22% | 53.67% | 48.63% | 41.13% | 27.23% |
| ProtT5 + Protein Top 2 | 59.23% | 53.82% | 48.47% | 40.91% | 27.37% |
| ProtT5 + Protein Top 3 | 59.15% | 53.63% | 48.10% | 41.03% | 27.44% |
| ProtT5 + Protein Top 4 | 59.19% | 53.84% | 48.34% | 41.26% | 27.53% |
| ProtT5 + Protein Top 5 | 58.99% | 54.00% | 48.51% | 40.89% | 27.11% |

*ProtT5 + Protein Top $n$* (where $n = 1$ to 5) indicates that the ProtT5 embeddings of the RBPs were combined with the $n$ protein sequence properties that registered the highest Gini importance after training the phage-host interaction prediction tool by Boeckaerts *et al.* [15] on our dataset. These sequence properties (in order of decreasing importance) are lysine frequency, isoelectric point, fourth protein Z-scale [57] (which is related to the heat of formation, hardness, electronegativity, and electrophilicity), percentage of residues with exposed solvent accessibility, and molecular weight. The header row refers to the confidence thresholds at which we evaluated model performance.

## Notes

### Competing Interest Statement

The authors have declared no competing interest.

